# A high-density genetic map and molecular sex-typing assay for gerbils

**DOI:** 10.1101/567016

**Authors:** Thomas D. Brekke, Sushmita Supriya, Megan Denver, Angharad Thom, Katherine A. Steele, John F. Mulley

## Abstract

We constructed a high-density genetic map in Mongolian gerbils (*Meriones unguiculatus*). We genotyped 137 F2 individuals with a genotype-by-sequencing (GBS) approach at over 10,000 loci and built the genetic map using a two-step approach. First, we chose the highest-quality set of 485 markers to construct a high-quality framework map of 1,239cM with 22 linkage groups as expected from the published karyotype. Second, we added an additional 5,449 markers onto the map based on their genotype similarity with the original markers. We used the final marker set to assemble 1,140 genomic scaffolds (containing ∼20% of annotated genes) into a chromosome-level framework. We used both genetic linkage and relative sequencing coverage in males and females to identify X- and Y-chromosome scaffolds and from these we designed a robust and internally-controlled PCR assay to determine sex. This assay will facilitate early stage sex-typing of embryonic and young gerbils which is not possible using current visual methods.

---

Rodents in the gerbil subfamily (Rodentia, muridae, gerbillinae) have been an important model for a huge range of organismal and evolutionary research. Gerbils inhabit the arid semi-deserts and steppes of Africa, Asia, and the Indian subcontinent and exhibit a wide range of adaptations to low water availability and poor food quality such as increased kidney function (Wilber and Gilchrist 1965), digestive function (Liu and Wang 2007) and altered insulin activity (Hargreaves et al. 2017). The extreme environmental pressures and a strong social structure have adapted Mongolian gerbil females to have a high degree of control over the sex ratio, even the ability to skew sex ratio in the left and right uterine horns independently (Clark et al. 1994). Cytological work on karyotype evolution suggests that genome stability in gerbils may be low and rearrangements common, even more so than in other rodents (Benazzou et al. 1982). Unusual patterns of meiosis are reported in both Mongolian gerbils and fat sandrats, where the sex chromosomes either do not pair (Ashley and Moses 1980), or do not recombine (la Fuente et al. 2007). Gerbils have also become popular as a medical model for a variety of human diseases. The great gerbil (*Rhombomys opimus*) is a known reservoir of human pathogens including the plague (Nilsson et al. 2018) and leishmaniasis (Ahmad 2002). Research on the great gerbil has spanned ecology (Linné Kausrud et al. 2007) and immune function (Nilsson et al.2018) in an effort to mitigate these major human health concerns. The fat sandrat (*Psammomys obsesus*), being well adapted to low calorie food sources, is highly susceptible to diabetes and is thought to share a similar genetic architecture as humans (Shafrir and Ziv 2009) which makes it an ideal model system. The Mongolian gerbil (Meriones unguiculatus) is also susceptible to diabetes (Li et al. 2016) and in addition has been a research model for epilepsy (Buckmaster 2006), stroke (Vincent and Rodrick 1979), and hearing loss (Abbas and Rivolta 2015).

While a great deal of cytological research has been done on various gerbils (Cohen 1970; Benazzou et al. 1982), there has been no bridge yet between the new genomics era and these classic karyotype studies. Three gerbil species (Mongolian gerbils, fat sandrats, and great gerbils) have all recently had their genomes sequenced (Hargreaves et al. 2017; Zorio et al. 2018; Nilsson et al. 2018). But as yet, none of these genomes are complete and all assemblies are highly fragmented. Many next-generation sequencing protocols include PCR steps where low GC-content regions amplify more efficiently than high GC-content regions (Tilak et al. 2018) This amplification bias has led to the omission of high-GC content genes in birds (Botero-Castro et al. 2017), and is likely responsible for missing regions in the gerbil genomes. One example is a missing region approximately five megabases long around the ParaHox cluster of Gsx1, Pdx1 and Cdx2. Due to its high GC-content a great deal of special effort to enrich the libraries for GC-rich DNA was required to successfully sequence the ParaHox region in fat sandrats (Hargreaves et al. 2017), and such effort has not been made for the other two gerbil species. Nor has any effort been made to increase contiguity of the gerbil genomes. Indeed, the least fragmented genome is from the great gerbil and contains only 6,390 scaffolds (Nilsson et al. 2018) which is impressive for shotgun sequencing, but not a chromosome-level assembly. In contrast, the fat sandrat genome has 150,763 scaffolds and the Mongolian gerbil genome has 68,793. Experiments looking into karyotypic evolution are stymied by so many small fragments. Constructing the fragmented genomes into chromosome-level assemblies will provide an important link between current large-scale sequencing projects and the classic cytological research on karyotype evolution and genomic rearrangements.

Despite having their genomes sequenced, gerbils still lack a published molecular sex-typing assay of the type that has been available for mouse for many years (Lavrovsky et al. 1998). Clark et al (1990; 1994) used visual inspection of the anogenital area to sex-type embryos in Mongolian gerbils and to test for sex-ratio skew *in utero*. They report an impressive 0% error rate but this takes considerable experience and a significant resource investment to check a subset of animals by marking them and allowing them to mature. One outcome of a chromosomal-level genome assembly is the ability to design a reliable and robust molecular sex-typing PCR assay. Such an assay would be less error-prone and available to a much wider community of gerbil researchers. Here we use an F2 mapping panel to construct a genetic map for Mongolian gerbils and use it to assemble scaffolds into chromosome-scale fragments. Based on sex-linkage we designed a robust and internally-controlled PCR assay to determine sex of gerbils.

## Methods

### Animal breeding and husbandry

Mongolian gerbils (*Meriones unguiculatus*) were housed at Bangor University under a 14 hour light to 10 hour dark daylight regimen and fed ab *libitum* in accordance with European Union and Home Office animal care regulations. All experiments were reviewed and approved by the Bangor University Animal Welfare and Ethical Review Board. A female from the Edinburgh strain and a male from the genetically distinct Sheffield strain (characterised in Brekke et al (2018)) were used as the parents of an F2 mapping panel. Four male and four female F1 individuals were crossed in pairs and produced 137 F2 offspring. All animals were euthanised using a Schedule 1 method and liver tissue from both parents, the eight F1s, and all 137 F2s was collected, snap-frozen in liquid nitrogen and stored at −80°C until use. Ear punches taken to identify animals during routine colony maintenance were placed directly into Buffer A from the MyTaq Extract-PCR kit (Bioline).

### DNA extraction and sequencing

Genomic DNA was extracted with a DNeasy Blood and Tissue DNA extraction kit (Qiagen) including the optional addition of 1 ul of 10mg/ml RNase added as per the manufacturer’s instructions. DNA was shipped to LGC Genomics (Queens Road, Teddington, TW11 0LY) for genotyping with a Genotyping-by-Sequencing (GBS) approach (Elshire et al. 2011) using the restriction enzyme MslI (recognition sequence: CAYNN^NNRTG). Overall genetic diversity is low between the Bangor and Edinburgh strains (Brekke et al. 2018) and so to identify sufficient variants we sequenced the 150bp paired-end libraries three times with two different size-selection regimes on three lanes of Illumina NextSeq 500 v2. The first two sequencing efforts were size-selected for reads 200-300 bases long while the final was selected for 300-380 bases. LGC Genomics demuliplexed the libraries, trimmed adapters, filtered reads for the presence of the MslI cut-site, and provided us with the resulting 770,140,572 total reads (5,239,051 reads per individual on average).

### Variant calling

Variant sites were identified and genotypes were called using the Stacks pipeline (version 2 beta) (Catchen et al. 2011; Catchen et al. 2013). We ran ‘process_radtags’ with the flags: -t 140, --disable_rad_check, --len_limit 140, -c, and -q. For ‘ustacks’ we used -m 3, -M 2, -N 4, -H, -d, --max_locus_stacks 3, --model_type snp, and --alpha 0.01. For ‘cstacks’ we used -n 2, and for ‘sstacks’, ‘tsv2bam’, and ‘gstacks’ we included the population map with -M. The population map included all individuals as a single population. The sequence of all stacks can be found in fasta format in Supplemental File 3. We ran ‘populations’ without the population map but included the flags -p 1, --min_maf 0.01, -- write_random_snp, and --vcf. Much segregating genetic diversity is shared between the two parental gerbil strains (Brekke et al. 2018) and so we identified 3,751 SNPs where the parents were homozygous for alternative alleles to use for building the genetic map. We also identified 7,063 SNPs that were heterozygous in one parent but not the other as some of these can be placed on the genetic map depending on which variant was inherited by each F1.

### Genetic map construction

We used r/QTL (Broman et al. 2003) to build the genetic map as described in Broman (2010). Two F2 individuals were removed from the panel due to low sequencing coverage. We filtered the 3,751 sites with variants that were homozygous for alternate alleles in the parents for ones that were genotyped in over 84% of F2 individuals and screened out ones with duplicate genotypes which resulted in a set of 485 high-quality markers. We used a LOD cutoff of 4 and a maximum recombination fraction of 0.30 to sort these markers into 51 linkage groups. These linkage groups were visually inspected for chromosome pairs with high LOD scores but low recombination fraction as this pattern indicates switched alleles. For each of these pairs, alleles were switched in one partner and the linkage groups were merged resulting in 22 chromosomes. Markers were ordered within their linkage groups with the function orderMarkers() and ripple(). The X was identified by looking for patterns of segregation distortion as X-linked markers will appear to show strong distortion under a Mendelian model of segregation of autosomes. To finalise the map, we sequentially dropped each marker to find ones that disproportionately expanded the map and removed these if they were not on the end of the chromosome. Using crossovers, we identified mistaken genotypes as ones that forced a double cross-over in a small distance and removed these. Finally we re-tested for segregation distortion while accounting for the hemizygosity of the X and found none. The script to run this is Meriones_Map_rqtl2.R and is available in the Supplemental File 5.

The 485 highest-quality markers were used to build the genetic map which left 10,329 genotyped SNPs, many of which could be associated with locations on the map. In order to incorporate the remaining SNPs, we removed uninformative genotypes for those loci from the dataset. Uninformative genotypes were those where the pattern of segregation of the alleles did not contain any information on the recombination that occurred between them.Specifically, when one parent was heterozygous at a site while the other was homozygous, the F1s can be either homozygous or heterozygous. If two heterozygous F1s were crossed, the F2 offspring genotypes are directly comparable with the genotypes of the original 485 markers used to build the genetic map and are therefore informative. Instead, if one or both F1s were homozygous at a site, then the genotypes at that site in their F2 progeny are not comparable to the SNPs used to build the map. Thus, we tracked the alleles at each potential locus through the known pedigree for each F2 individual to identify and remove uninformative genotypes.

The genotypes of the remaining informative loci were compared with the genotypes of all mapped loci for every individual and we counted the number of matching genotypes to identify the most similar marker on the map. Each new marker was associated with the map if it shared greater than 90% of the genotypes with any marker. New markers were placed at the centiMorgan location of the first marker with which it shared the most similar genotypes (Supplemental Figure 1). We did not allow markers added in this way to change the order nor spacing of the original map.

**Figure 1:**
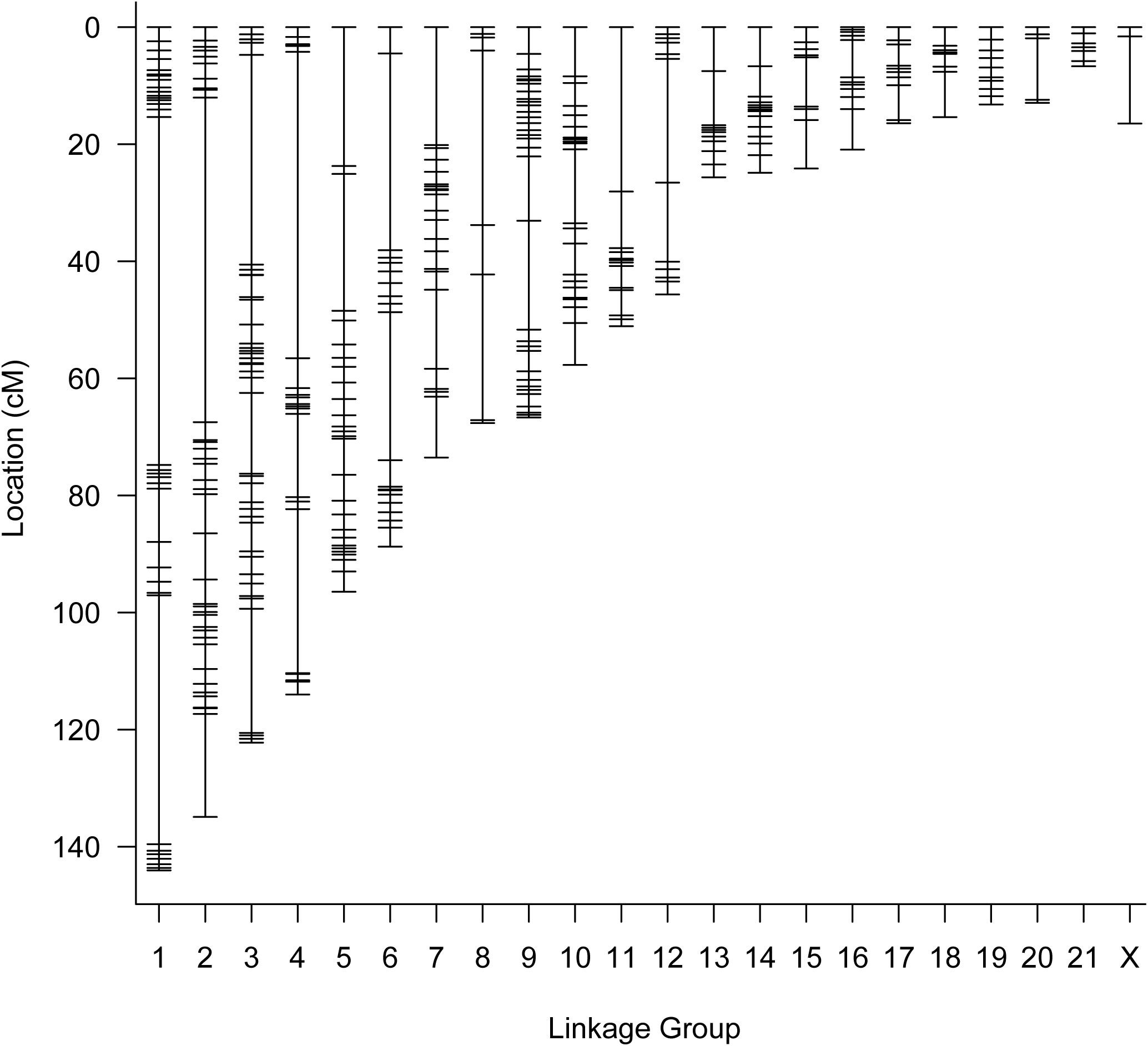
Genetic map of the Mongolian gerbil genome. The map contains 6034 SNP markers, assigned to 21 autosomes and the X chromosome, and comprises 1239.1cM.

### Scaffolding the genome

We aligned the raw GBS reads to the reference stacks (Supplemental File 3) with bwa (Li and Durbin 2009) and calculated the depth of coverage for each individual with samtools (Li et al. 2009). By parsing the depth of coverage by sex we identified sex-linked markers as described in (Brekke et al. 2018) based on the logic that markers on the Y have coverage in males but not females, scaffolds on the X have twice the coverage in females as in males, and autosomal markers have approximately equal coverage in males and females. To calculate the standardised average coverage for males and females, we divided all read counts for each individual by the sequencing effort of that individual, multiplied by 1,000,000, and then took the mean of all males and all females. Y linked markers fulfil the inequality:

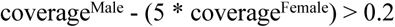

Unknown markers are ones that are not Y-linked and where:

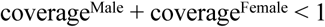

X-linked markers are neither unknown nor Y-linked, and satisfy the inequality:

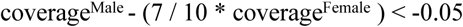

All other markers are autosomal. The regions demarcated by these cutoffs are shown in detail in Supplemental Figure 2. Then we used blastn to align each marker to the gerbil genome GCF_002204375.1_MunDraft-v1.0_genomic.fna (Zorio et al. 2018). Once markers were aligned to the genome we annotated sex-linked scaffolds and tested how often a single genomic scaffold associated with multiple different linkage groups as these cases indicate chimeric scaffolds. Chimeric scaffolds were removed from further analysis. Finally, we calculated how many scaffolds and bases were associated with the genetic map.

**Figure 2:**
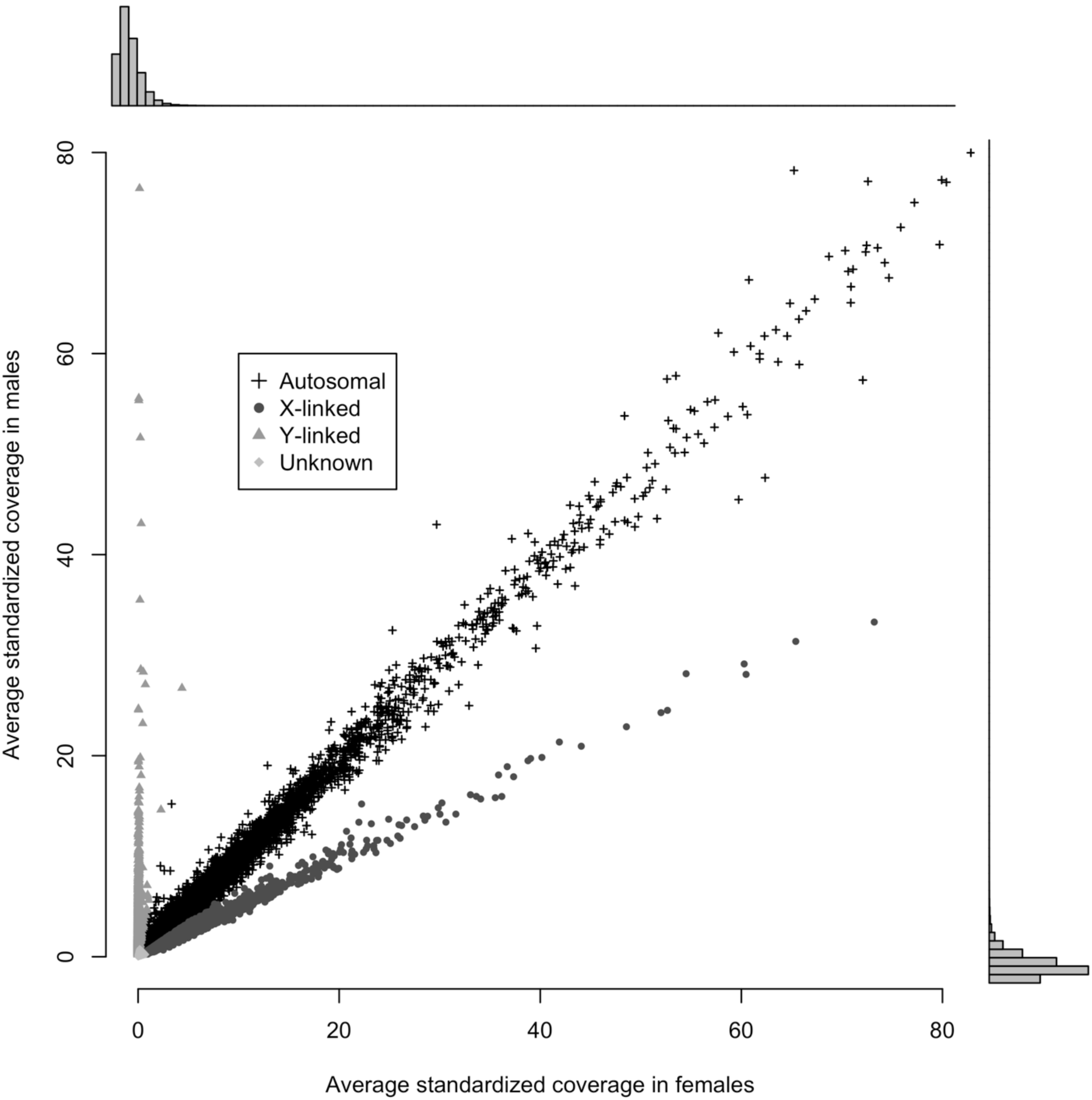
Sex-specific coverage of markers. Y-linked markers have no coverage in females, and X-linked markers have twice the coverage in females as males.

**Figure 3:**
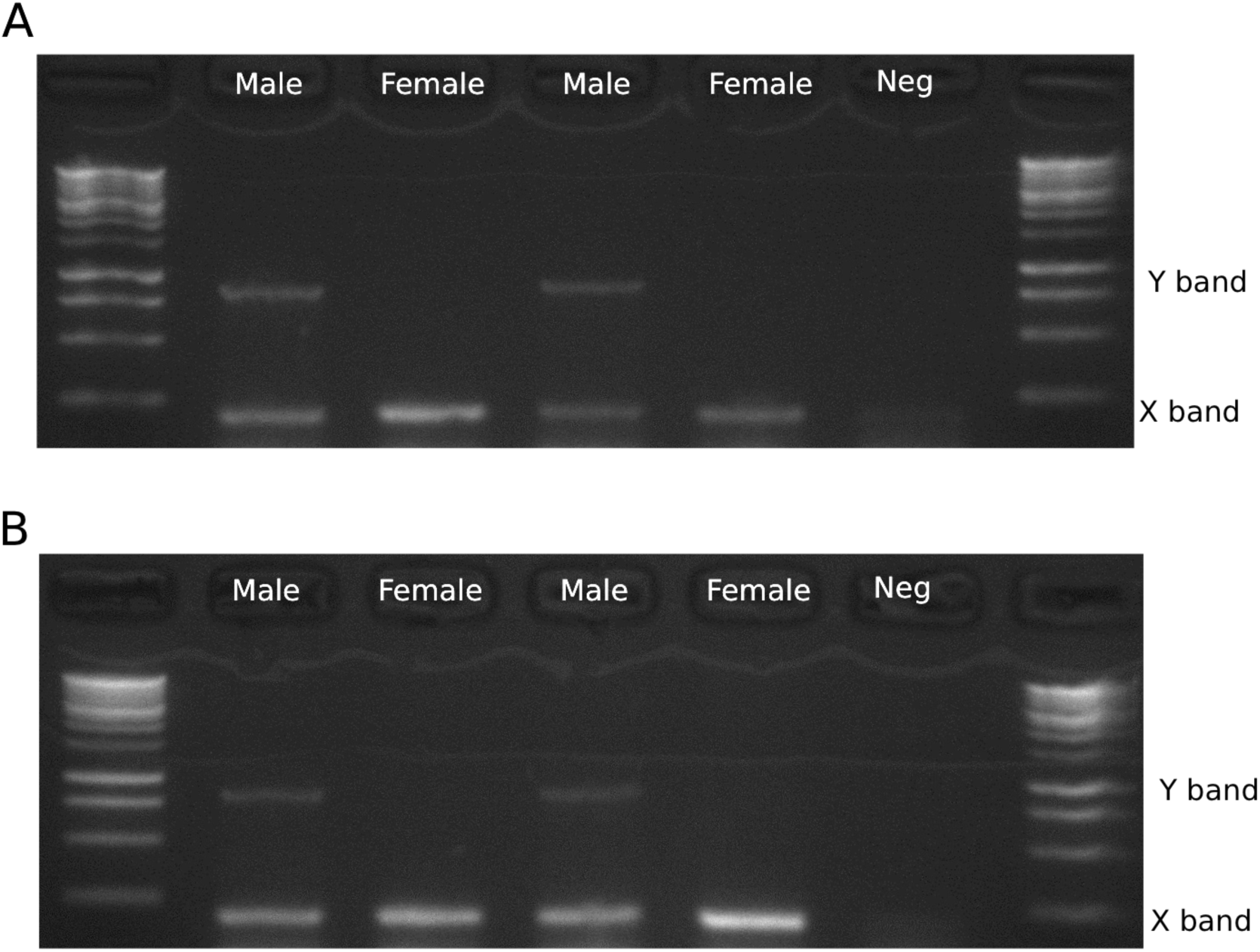
PCR assay for sex-typing gerbils. DNA extracted from liver for both Mongolian gerbils (A) and fat sandrats (B) can be robustly amplified with the primer pair. Males amplify two bands while females amplify one. The Y-linked band is at 845 bases while the X-linked band is at 206 bp. The ladder is the Bioline 1kb ladder. ‘Neg’ is the no-template control.

### Sex-typing assay

We chose two sex-linked scaffolds from our annotation: one on the X chromosome (NW_018661451.1) and one on the Y (NW_018662972.1). We used Primer3 (Untergasser et al. 2012; Koressaar et al.) to design PCR primers to each of these scaffolds. Primer pairs were chosen such that the X and Y bands would be easily distinguishable based on product length, with similar melting temperatures, and little complementarity so that they could be run in the same reaction. We optimised the multiplex PCR reaction under a range of temperature and cycle number conditions in both males and females to find a set where the X-linked primers amplified in both males and females while the Y-linked primers only amplified in males. A multiplexed two-step PCR worked best with the following conditions: an initial denaturation at 95°C for 1 minute, followed by 30 cycles of 94°C for 15 seconds and 72°C for 1 minute with a final extension step of 72°C for 5 minutes. We tested the primers on DNA extracted from the liver of Mongolian gerbil and fat sandrat using the MyTaq Red Mix (Bioline). In each 25ul PCR reaction, we used 100ng of DNA (2ul of 50ng/ul) and 0.5ul of each of the four 10nM primers.

## Results

### Genetic map

We created a high-quality genetic map for Mongolian gerbils with 21 autosomal linkage groups and the X. This map spans 1239.1cM and includes 485 SNP markers (Figure 1, Table 1, Supplemental File 1). Onto this framework we were able to place an additional 5,549 SNP markers based on genotype similarity (Supplemental Figure 1, Supplemental File 2).

**Table 1:** Map statistics for the Mongolian gerbil genetic map

| Linkage group | Length in cM | Number of markers<br>in base map | Average marker<br>spacing (cM) | Max marker<br>spacing (cM) | Number of markers<br>in final map |
| --- | --- | --- | --- | --- | --- |
| **1** | 144.0 | 44 | 3.3 | 59.4 | 616 |
| **2** | 134.9 | 56 | 2.5 | 55.4 | 689 |
| **3** | 122.2 | 51 | 2.4 | 35.8 | 540 |
| **4** | 114.0 | 28 | 4.2 | 52.3 | 448 |
| **5** | 96.4 | 28 | 3.6 | 23.7 | 402 |
| **6** | 88.7 | 22 | 4.2 | 33.6 | 333 |
| **7** | 73.5 | 26 | 2.9 | 20.1 | 333 |
| **8** | 67.6 | 8 | 9.7 | 29.8 | 152 |
| **9** | 66.7 | 42 | 1.6 | 18.6 | 481 |
| **10** | 57.7 | 34 | 1.7 | 12.6 | 304 |
| **11** | 51.1 | 21 | 2.6 | 28.1 | 195 |
| **12** | 45.7 | 13 | 3.8 | 21.1 | 226 |
| **13** | 25.6 | 13 | 2.1 | 9.3 | 109 |
| **14** | 24.9 | 19 | 1.4 | 6.7 | 211 |
| **15** | 24.1 | 12 | 2.2 | 8.4 | 200 |
| **16** | 20.9 | 13 | 1.7 | 6.9 | 151 |
| **17** | 16.4 | 14 | 1.3 | 6 | 205 |
| **18** | 15.4 | 13 | 1.3 | 7.7 | 125 |
| **19** | 13.2 | 11 | 1.3 | 2.1 | 114 |
| **20** | 12.9 | 7 | 2.2 | 10.5 | 82 |
| **21** | 6.7 | 7 | 1.1 | 1.7 | 56 |
| **X** | 16.5 | 3 | 8.2 | 14.9 | 62 |
| **Overall** | 1239.1 | 485 | 2.7 | 59.4 | 6034 |

We annotated 421,257 autosomal stacks, 11,322 X-linked stacks, 5,110 Y-linked stacks, and 14,505 unknown stacks by analysing the relative coverage of males and females (Figure 2). By aligning these stacks to the published genome (Zorio et al. 2018), we annotated 17,121 autosomal scaffolds (2,246,217,465 base-pairs), 3,766 X-linked scaffolds (322,528,372 base pairs), and 2,158 Y-linked scaffolds (71,814,401 base pairs) (Supplemental File 4).

### Scaffolding the genome

Using information from the genetic map position of SNP markers and their alignments to the published genomic scaffolds, we were able to place 1,140 scaffolds onto the chromosomal framework. In addition, we identified 12 putatively chimeric scaffolds which were removed from further analysis. The 1,140 scaffolds span 400,346,323 sequenced bases (15.9% of bases) and include 4,603 genes (19.8% of annotated genes).

### Sex-typing

A robust sex-typing PCR test was designed to use primer pairs that act as two molecular markers that can be multiplexed, one marker for each for the X and Y chromosomes (Table 2). The X-linked primers amplified a 202 base-pair fragment in the gene *Kdm5c* and the Y-linked primers amplified a 845 base-pair fragment in the gene *Uba1Y*. Fragments from both the X and Y markers were present in all males while the Y primers did not amplify in females (Figure 2). This PCR assay amplified DNA extracted from liver in both Mongolian gerbils and fat sandrats.

**Table 2:**
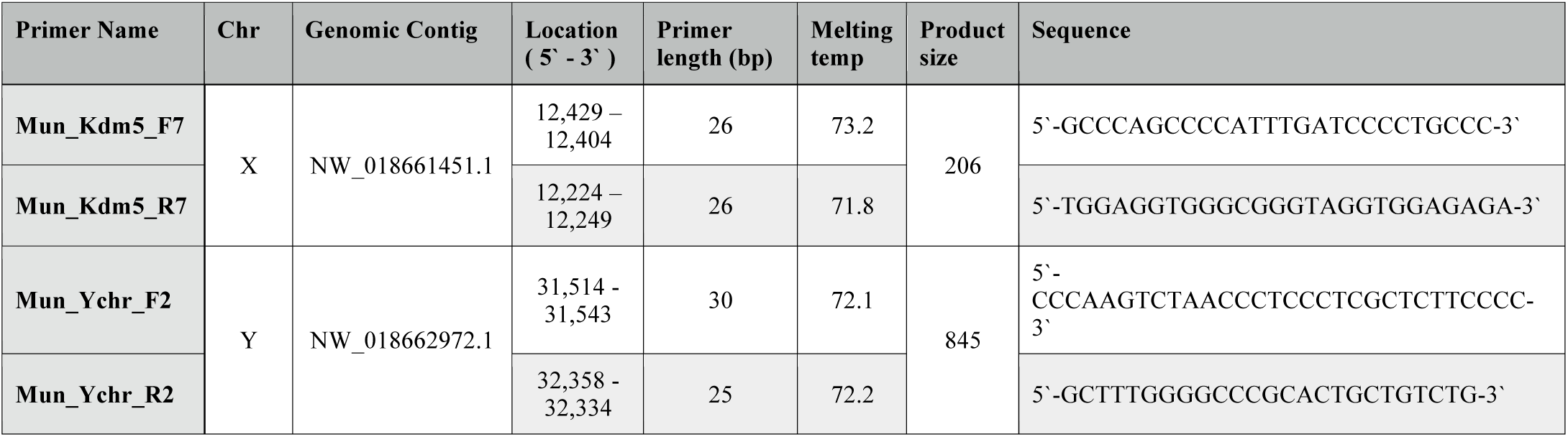
Primer details for the X and Y chromosome-specific PCR-based sex assay.

## Discussion

After years of being side-lined by geneticists in favour of other model mammals, gerbils and their relatives are at last entering the genomic era and, with the advent of new technologies and massive reductions in sequencing costs, it seems likely that we will see multiple chromosome-scale assemblies for this group in the near future. Our analysis is a first step towards tying genomic scaffolds to chromosomes, and will hopefully pave the way for future work aimed at both improving the assembly of the Mongolian gerbil genome, and facilitating cross-species comparisons, particularly for the reconstruction of patterns of chromosomal evolution

In order to measure recombination frequency and build the map it is easiest to use markers that have variants alternatively homozygous in the parents. To maximise the number of these markers available for mapping we used a cross between the two most divergent strains of gerbil locally available: Edinburgh and Sheffield (Brekke et al. 2018). These strains have been bred in isolation for a number of years but do ultimately originate from the same Tumblebrook Farm strain stock (Brekke et al. 2018). While some variation has fixed within each strain, the majority of variation segregates in all strains. To account for this heterozygosity we screened for variants that were homozygous for different alleles in the parents and used these for map building, later adding on other informative sites to the backbone of the map.

Until recently, the limiting factor in genetic mapping experiments was identifying the markers. Maps would commonly consist of a couple hundred markers. For instance in 1992 a genetic map for house mice included 317 markers (Dietrich et al. 1992) and the first map for rats had 432 (Jacob et al.1995). Developing and genotyping markers was enormously expensive compared to the cost of breeding and raising the animals needed for the mapping panel. Thus the number of animals, and hence the amount of recombination events available to discriminate marker location was often well in excess of what was necessary for the number of markers on the map. Today, next-generation sequencing technology allows us to discover millions of markers with very little effort or expense and breeding a mapping population large enough to accurately order thousands to millions of markers can be hugely expensive. We are now in era of recombination-limitation instead of marker-limitation, and run the risk of creating high-density maps that are not reliable at the fine-scale. To circumvent this issue, researchers have recently begun to identify a subset of markers to create a high-quality map using few enough markers that match the number of recombinations in the mapping panel. Onto these base maps are layered as many additional markers as possible, either by grouping markers into “0-recombinant clusters” as in (Li et al. 2015) or by using a regression algorithm to layer additional markers onto the map as in (Blankers et al. 2019). Here we use a simple genotype matching algorithm that identifies groups of identical or near-identical markers with which to associate each additional marker. All these approaches result in a high density genetic map, but make explicitly clear that the fine-scale order of the additional markers is unknowable given the size of the mapping panel.

The mapping panel we used contained 137 individuals, so we selected 485 markers that had the least missing data with the aim of finding 10-20 markers per chromosome which is enough to build a linkage group, but not so many that would saturate the number of recombination events present. Using these markers, we built a genetic map containing 22 linkage groups, the same as the number of chromosomes in the Mongolian gerbil genome (Cohen 1970). Onto this high-quality map we added in as many additional markers as possible. These include the markers alternatively homozygous in the parents as well as many markers that were heterozygous in one parent whose inheritance we traced through the F1s. New markers were associated with a specific location on the map but not allowed to alter its length nor the order of the original high-quality markers.

In addition to genetic linkage, the sex chromosomes can be annotated using the relative coverage of males and females. With this method, we have annotated 3,766 X-linked scaffolds, and 2,158 Y-linked scaffolds. Together these scaffolds account for 322,528,372 bases on the X chromosomes and 71,814,401 bases of the Y. This approach does not provide any genomic order for these scaffolds, but it allowed us to design a reliable molecular sex-typing assay for gerbils. Our sex-typing assay relies on two primer pairs that simultaneously amplify X- and Y-linked regions in the same PCR reaction. Thus each reaction is internally controlled and does not rely on the failure of PCR to positively ID a female, unlike the SRY-based approach taken in a variety of species (Shaw et al. 2003; Kusahara et al. 2006; Prashant et al. 2008). Indeed, PCR failure is easily identified due to the absence of all bands and so this approach mitigates errors where males are incorrectly identified as female. A molecular sex-typing approach opens the door to a variety of novel experiments and that require sex-typing embryonic and juvenile individuals. Sex-typing juvenile animals using a non-lethal approach (such as with DNA extracted from an ear-punch or toe-clip) will greatly reduce the daily costs associated with rearing and maintaining animals that are not needed for a sex-specific experiment or routine colony maintenance. Additionally, a molecular assay paves the way for a large-scale analysis of embryonic sex ratio bias in Mongolian gerbils.

## Supporting information

Supplemental_File_1_Meriones_Genetic_Map

Supplemental_File_2_Meriones_Genetic_Map_allMarkers

Supplemental_File_3_Stacks_catalog.fasta

Supplemental_File_4_Marker_annotation_and_scaffold

Supplemental_File_5_Codebase

## Acknowledgements

We would like to thank the animal care technicians at Bangor University: Rhys Morgan, Rebecca Snell, Mike Hayle, and Emlyn Roberts without whose oversight, care, and guidance our animal work would not have been possible. We would also like to thank Kris Crandell, Alex Papadopulos, Gil Smith, and Mark Quinton-Tulloch for helpful discussions throughout the duration of this project. This research was funded by a Leverhulme Trust research project grant to J. F. M. and K. A. S. (RPG-2015-450).

## Figures

**Supplemental Figure 1:**
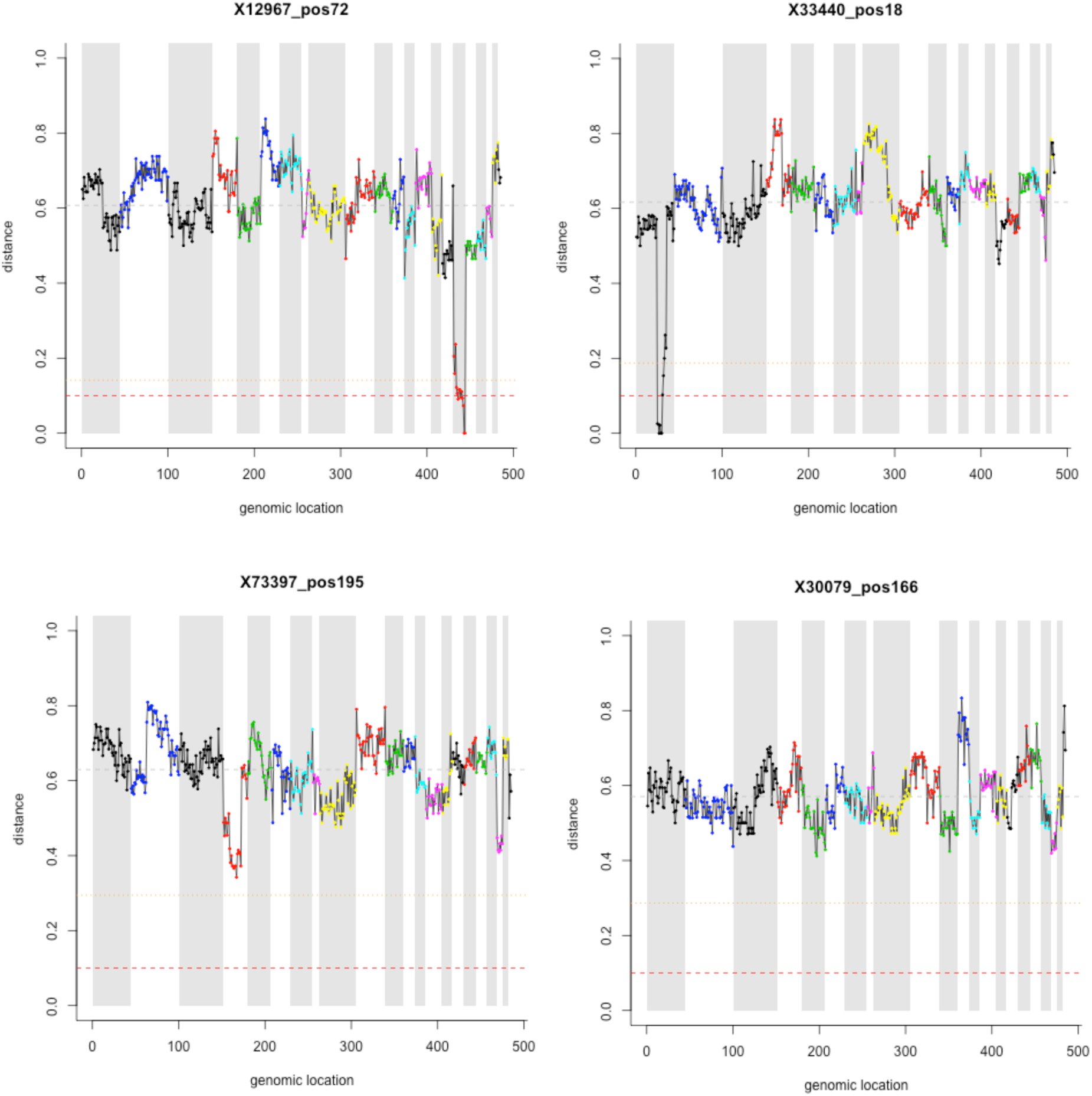
Placing additional markers on the map. Two examples of markers with strong association at specific locations on the map (top row: 12967_pos72 and 33440_pos18) and two examples of markers without a strong association (bottom row: 73397_pos195 and 30079_pos166). The X-axis is the ordered set of the 485 high-quality markers that comprise the map. Linkage groups are differentiated by point colour and alternating grey background stripes. The Y-axis is the genotype distance between two markers, measured as the proportion of mismatching genotypes in the comparison between the unplaced marker and each map location. The grey dashed line is the genome-wide average, the orange dotted line is 4 standard deviations away from the mean, and the red dashed line is the 10% cutoff. Markers were only associated with the map if they had fewer that 10% mismatching genotypes, in which case they were placed at the same centiMorgans as the first marker with which they shared the most matching genotypes.

**Supplemental Figure 2:**
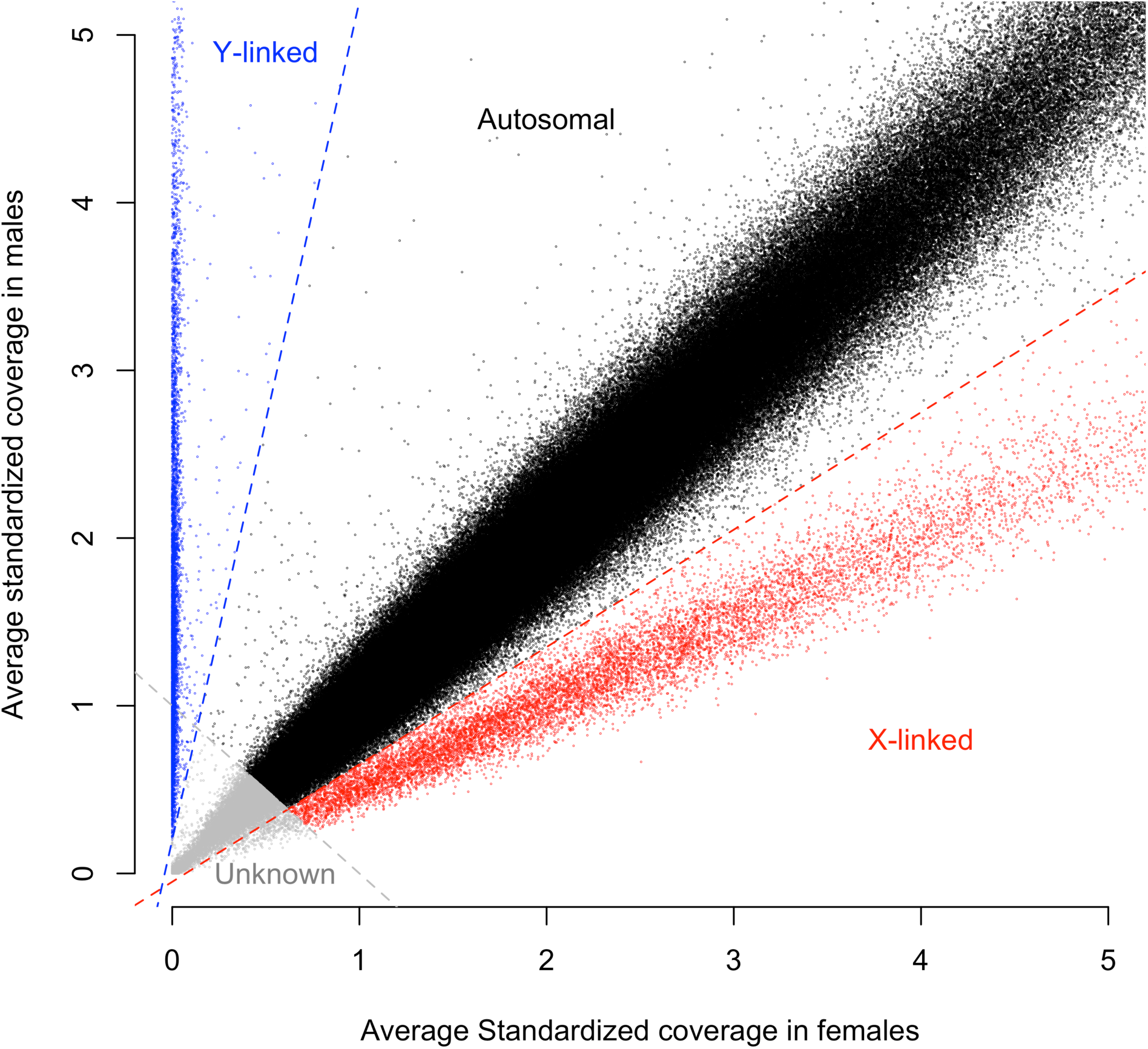
Supplemental Figure 2: Boundaries for annotating sex-linkage in markers. This shows an extreme close-up of the cutoffs chosen to differentiate Autosomal, X, and Y linked markers. The blue line distinguishing Y-linked markers is at: coverage^Male^ = (5 * coverage^Female^) + 0.2, the grey line to identify unknown markers is at coverage^Male^ = 1 - coverage^Female^, and the red line to distinguish X-linked markers is coverage^Male^ = (7 / 10 * coverage^Female^) - 0.05. These cutoffs were chosen by visually inspecting this plot for the natural breakpoints between Autosomal-, X-, and Y-linked makers with the observation that Y-linked markers cluster at coverage^Female^ = 0, Autosomal markers cluster at coverage^Male^ = coverage^Female^, and X-linked markers cluster at coverageMale = 1/2 * coverage^Female^. The boundary for the unknown markers is set conservatively to avoid mistaking low-coverage X-linked and Autosomal markers.

Supplemental File 1: Meriones Genetic Map. The genetic map file in r/QTLs “csvr” format which includes the 485 original markers and genotypes of F2 individuals. Marker names are formatted to include the marker ID and the position of the SNP within the marker separated by an underscore as such: “168795_pos129”.

Supplemental File 2: Meriones Genetic Map with all Markers. The genetic map in r/QTLs “csvr” format including all 6,034 markers and genotypes. Uninformative genotypes have been removed. Marker names are formatted to include the marker ID and the position of the SNP within the marker separated by an underscore as such: “168795_pos129”.

Supplemental File 3: Stacks catalog fasta. A fasta file containing all stacks in the catalog created by the stacks pipeline.

Supplemental File 4: A list of scaffolds and which are autosomal, sex-linked, or unknown, and their location on the genetic map if known. This is a tab-separated file with five columns: marker, annotation, scaffold, linkage group, centiMorgans.‘Annotation’ can be A, X, Y, or U for autosomal, X-linked, Y-linked, or Unknown and was determined based on relative coverage in males and females. A “*” in the scaffold column indicates that the marker did not align to any scaffold.

Supplemental File 5: The code base for this project. All code can be run with “Meriones_Stacks_5.sh” which calls the other scripts. This code will require many dependent programs and packages to be installed such as stacks and r/QTL.

