## Supplemental_File_5_Codebase for "A high-density genetic map and molecular sex-typing assay for gerbils": add_het_markers_to_Meriones_Map.py

```
#!/usr/bin/env Python3
#add both het and hom markers - easier than 2 scripts - though the name is confusing. 

import sys
import re
#Grab het markers that are on the list called hetMarkers_placed.txt from csv: rqtl_input_Var_2.csvr and use the PandF1 dataset to modify the genotypes, then print them out and then add them to the Meriones Map 
def parse_het():
	with open("het_Markers_placed.txt", 'r') as HET:
		counter=0
		for line in HET:
			counter+=1
			line=line.strip("\n")
			line=line.lstrip("X")
			fields=line.split("	")
			#print(fields)	
			#print(fields[0:4], file=sys.stderr)
			rad=fields[0]	
			if rad not in het_dict:
				het_dict[rad]=",".join(fields[1:3])	
				het_dict2[rad]=fields[3]
			#else:
			#	print("	already present", file=sys.stderr)
	#print(het_dict, file=sys.stderr)			
	print("Read", counter, "lines of het_Markers_placed.txt. Added markers to het_dict as RAD_SNP=LG,pos. Example of RAD_SNP:", rad, "Example of LG,pos:", ",".join(fields[1:3]), file=sys.stderr)
def parse_hom():
	with open("hom_Markers_placed.txt", 'r') as HOM:
		counter=0
		counter2=0
		for line in HOM:
			counter+=1
			line=line.strip("\n")
			line=line.lstrip("X")
			fields=line.split("	")
			rad=fields[0]
			if rad not in hom_dict:
				counter2+=1
				hom_dict[rad]=",".join(fields[1:3])
				hom_dict2[rad] = fields[3]
	#print(hom_dict, file=sys.stderr)			
	print("Read", counter, "lines of hom_Markers_placed.txt.", counter2, "lines added to hom_dict.", file=sys.stderr)


#this one gives just the radtag ID
def parse_name(name):
	bits=name.split("_")
	if len(bits)==2:
		return(bits[0])
	elif len(bits)==6:
		return(bits[4])
	else:
		print("Error - name is odd", name, file=sys.stderr)	 		
		return("")

#this one gives the radtag ID AND the SNP position. 
def parse_name2(name):
	bits=name.split("_")
	if len(bits)==2:
		#print(name, bits, bits[0]+"_pos"+bits[1], file=sys.stderr )
		return(bits[0]+"_pos"+bits[1])
	elif len(bits)==6:
		return(bits[4]+"_pos"+bits[5])
	else:
		print("Error - name is odd", name, file=sys.stderr)	 		
		return("")
				
def parse_geno(geno):
	fields=geno.split(":")
	#print(geno)#GT:DP:AD:GQ:GL
	geno=fields[0]
	if geno == "./.":
		AD="0,0"
		GD=0
		GQ=0
		PL="0,0,0"
	else:
		AD=fields[2]
		GD=int(fields[1])
		GQ=int(fields[3])
		PL=fields[4]
	if GQ < 0:									#here we can pick at cutoffs
		geno="./."
	elif GD < 0:
		geno="./."	
	#print(geno, file=sys.stderr)	
	g=format_geno(geno)
	return (g)

def format_geno(f):
	if f=="0/0" or f=="BB":
		g="B"		
	elif f=="0/1" or f=="AB" or f=="BA":
		g="H"
	elif f=="1/1" or f=="AA":
		g="A"
	elif f=="./." or f=="-":
		g="N"
	else:
		print("The genotypes cannot be formatted correctly:", f, file=sys.stderr)	
	return(g)

def format_geno2(f):
	if f=="0/0" or f=="BB" or f=="B":
		g="BB"		
	elif f=="0/1" or f=="AB" or f=="BA" or f == "H":
		g="AB"
	elif f=="1/1" or f=="AA" or f=="A":
		g="AA"
	elif f=="./." or f=="-" or f=="NA":
		g="-"
	else:
		print("The genotypes cannot be formatted correctly:", f, file=sys.stderr)	
	return(g)

def get_cross_number(ID):
	#needs to take in the position in the genotypes string ie. 0 or 1 or 137 and return C1, C2, C3, or C4. 
	
	#I dropped two individuals from the panel - they are genotyped in the VCF, but not in the map. So this line is slightly different between add_het_markers_to_Meriones_map (which works with things after I dropped those individuals - so len=135 ) and make_rqtl_input_file_For_Capture which deals with the VCF (so len=137). 
	cross_origins=['C1', 'C1', 'C1', 'C1', 'C1', 'C1', 'C1', 'C1', 'C1', 'C1', 'C1', 'C4', 'C4', 'C4', 'C4', 'C4', 'C4', 'C2', 'C2', 'C2', 'C2', 'C2', 'C2', 'C3', 'C3', 'C3', 'C3', 'C3', 'C1', 'C1', 'C1', 'C1', 'C1', 'C1', 'C1', 'C1', 'C1', 'C1', 'C2', 'C2', 'C2', 'C2', 'C2', 'C2', 'C2', 'C4', 'C4', 'C4', 'C4', 'C4', 'C4', 'C4', 'C4', 'C3', 'C3', 'C3', 'C3', 'C3', 'C3', 'C3', 'C1', 'C1', 'C1', 'C1', 'C1', 'C1', 'C1', 'C4', 'C4', 'C4', 'C4', 'C4', 'C4', 'C4', 'C4', 'C4', 'C3', 'C3', 'C3', 'C3', 'C3', 'C1', 'C1', 'C1', 'C1', 'C1', 'C1', 'C1', 'C3', 'C3', 'C3', 'C3', 'C3', 'C3', 'C2', 'C2', 'C2', 'C2', 'C2', 'C2', 'C2', 'C2', 'C2', 'C2', 'C2', 'C2', 'C2', 'C2', 'C4', 'C4', 'C4', 'C3', 'C3', 'C3', 'C3', 'C3', 'C4', 'C4', 'C4', 'C1', 'C1', 'C1', 'C1', 'C1', 'C1', 'C1', 'C3', 'C3', 'C3', 'C3', 'C2', 'C2', 'C4', 'C4', 'C4', 'C4', 'C4']
	return(cross_origins[ID])

def get_c(geno1,geno2):
	C=geno1+geno2
	if re.search("N", C):#ungenotyped parent
		out=0
	elif C =="HH": #best - both F1s are hets
		out=2
	else: #one F1 is a het, the other is homozygous - this is possibly useable down the road. Ignoring it now.
		out=1
	return(out)

def get_useful_crosses(geneID,genos):
	#print(genos)
	P=genos[0]+genos[9]
	C1=get_c(genos[1],genos[4])
	C2=get_c(genos[2],genos[3])
	C3=get_c(genos[5],genos[8])
	C4=get_c(genos[6],genos[7])	

	if geneID not in useful_dict:
		useful_dict[geneID]={}
	useful_dict[geneID]["C1"]=C1 
	useful_dict[geneID]["C2"]=C2
	useful_dict[geneID]["C3"]=C3
	useful_dict[geneID]["C4"]=C4
	#now useful_dict can calculate back to whether a cross will have useful offspring -> 2=all offspring are useable!, 1=offspring mimic a backcross and probably only the hets should be used, 0=no offspring are usable from this cross. 

def parse_PandF1():
	with open("PandF1.recode.vcf", 'r') as VCF:
		for line in VCF:
			line=line.strip('\n')
			if line.startswith("#"):
				if not line.startswith("##"):
					#format headers here? Maybe extract individual ids?
					a=1
					#print(line, file=sys.stderr)
			else:	
				fields=line.split("	")
				gene_ID="_pos".join(fields[0:2])
				genotypes=fields[9:]
				genos=[parse_geno(genotype) for genotype in genotypes]
				get_useful_crosses(gene_ID, genos)
				#print(genos, useful)

#this one goes through and puts NA's in where the genotypes are uninformative. 
def recalculate_geno_VAR(line):
	fields=line.split(",")
	name=parse_name2(fields[0])
	genotypes=fields[2:]
	genos=[]
	for genotype,i in zip(genotypes,range(0,len(genotypes))):
			if useful_dict[name][get_cross_number(i)]==2:
				genos.append(format_geno2(genotype))
			else:
				genos.append("-")		
	return(genos)
	
def recalculate_geno_Fixed(line):
	fields=line.split(",")
	name=parse_name2(fields[0])
	genotypes=fields[2:]
	genos=[format_geno2(genotype) for genotype in genotypes]
	#print(genos)
	return(genos)
		
def drop_bad_inds(fields):
	#"ESES1-5M" and "ESES14-7F" were dropped form the map and shouldn't be included. 
	#print(len(fields), end="	")	
	del fields[35]  #index 36 is individual ESES14-7F - python base 0 means index 35
	del fields[4]	#index 5 is individual ESES1-5M   - python base 0 means index 4
	#print(len(fields), end="\n")
	return(fields)

def parse_rqtl_input_Var():
	with open("rqtl_input_Var_2.csvr", 'r') as RQTL:
		counter=0
		counter2=0
		for line in RQTL:			
			counter+=1
			if counter>7:
				line=line.strip("\n")
				fields=line.split(",")
				name=fields[0]
				rad=parse_name2(name)
				if rad in het_dict: # and rad not in already_in_there_dict:
					counter2+=1
					genos=recalculate_geno_VAR(line)
					genos=drop_bad_inds(genos)
					if het_dict2[rad] == "YES":
						print(rad, file=sys.stderr)
						print(genos, file=sys.stderr)
						genos = switch_alleles(genos)
						print(genos, file=sys.stderr)

					print(name, het_dict[rad], ",".join(genos), sep=",", end="\n" )
					############################
				#	if rad not in printed_hets:#troubleshooting - can hash out
				#		printed_hets[rad]=0    #
				#	printed_hets[rad]+=1	   #
					############################					
		print("Read", counter-7, "lines of rqtl_input_Var_2.csvr. Checked if Rad was in het_dict, if so formatted the genotypes and printed the output.Example rad is:", rad, "\n\t", counter2, "lines printed.", file=sys.stderr)

def parse_rqtl_input_Fixed():
	with open("rqtl_input_2all.csvr", 'r') as RQTL:
		counter=0
		counter2=0
		for line in RQTL:			
			counter+=1
			if counter>7:
				line=line.strip("\n")
				fields=line.split(",")
				name=fields[0]
				rad=parse_name2(name)
				#print(rad)
				if rad in hom_dict and rad not in already_in_there_dict:
					counter2+=1
					genos=recalculate_geno_Fixed(line)
					genos=drop_bad_inds(genos)
					if hom_dict2[rad] == "YES":
						print(rad, file=sys.stderr)
						print(genos, file=sys.stderr)
						genos = switch_alleles(genos)
						print(genos, file=sys.stderr, end="\n\n")
					print(name, hom_dict[rad], ",".join(genos), sep=",", end="\n" )
					############################
					#if rad not in printed_homs:#this is for troubleshooting - can hash out
					#	printed_homs[rad]=0    #
					#printed_homs[rad]+=1	   #
					############################
			
				#else:
					#print(rad, file=sys.stderr)	
		print("Read", counter-7, "lines of rqtl_input_2.csvr. Checked if Rad was in hom_dict and not in already_present_dict, if satisfied, formatted and printed genotypes.\n\t", counter2, "lines were printed which is all the lines read less the markers that made up the backbone of the map.", file=sys.stderr)

def switch_alleles(genos):
	switch_dict={"AB":"AB", "AA":"BB", "BB":"AA", "-":"-"}
	new_genos = [switch_dict[g] for g in genos]
	return(new_genos)
		
def parse_map():
	with open('Meriones_Map.csv') as MAP:
		counter=0
		for line in MAP:
			counter+=1
			if counter>7:
				name=line.split(",")[0]
				rad=parse_name2(name)
				if rad not in already_in_there_dict:
					already_in_there_dict[rad]=0
				already_in_there_dict[rad]+=1		
	print("Read", counter-7, "lines of Meriones_Map.csv. Added markers to already_in_there_dict as: RAD=count_of_Rad", file=sys.stderr)


useful_dict = {} # shows which tags work for which crosses
hom_dict = {}
hom_dict2 = {}
het_dict = {}	#lists the het markers that are worth placing	
het_dict2 = {}
already_in_there_dict = {}
printed_hets = {}
printed_homs = {}

parse_map()
parse_het()
parse_PandF1()
parse_rqtl_input_Var()
parse_hom()
parse_rqtl_input_Fixed()

#print(already_in_there_dict, file=sys.stderr)


#print("\nMarkers that did not appear in the printed_homs (from rqtl_input_var_2), but did appear in the het_dict (from het_Markers_placed)", file=sys.stderr)
#for rad in hom_dict:
#	if rad not in printed_homs:
#		print(rad, hom_dict[rad], file=sys.stderr)		
#print("Here is the average number of entries in printed_hets (should be 1, not more):", sum([float(printed_hets[i]) for i in printed_hets])/len(printed_hets), file=sys.stderr)
#for i in useful_dict:
#	print(i, useful_dict[i])

#for i in het_dict:
#	print(i, het_dict[i])
```
