## Supplemental_File_5_Codebase for "A high-density genetic map and molecular sex-typing assay for gerbils": Annotate_AXY_Meriones_2.R

```
#!/usr/local/bin/Rscript

#a script to annotate the sex chromosomes of the gerbils. 
#use for the Mapping panel - not the GBS diversity paper

#for HPC wales
#setwd("/scratch/thomas.brekke/Meriones_GBS/alignments/")
#cov<-read.table("coverage_repDepth.txt", stringsAsFactors=F, header=T)

#for testing the script on Mwk16:
#setwd("/Volumes/university/College of Natural Sciences/Research Data/BIOL-GROUP-Mulley/tom_brekke/Meriones_GBS/alignments/")
setwd("/Volumes/LaCie/Projects/Meriones_Map/alignments/")
covFILE="coverage_repDepth.txt"
covb<-as.data.frame(read.table(covFILE, stringsAsFactors=F, header=F));Names<-c("chr", "pos","EEEE0-1F", "EESS2-1F", "EESS2-3F", "EESS2-4M", "EESS2-6M", "EESS3-1F", "EESS3-2F", "EESS3-3M", "EESS3-4M", "ESES1-1F", "ESES1-2M", "ESES1-3M", "ESES1-4M", "ESES1-5M", "ESES1-6M", "ESES10-1M", "ESES10-2F", "ESES10-3M", "ESES10-4F", "ESES10-5F", "ESES11-1F", "ESES11-2M", "ESES11-3", "ESES11-4", "ESES11-5", "ESES11-6", "ESES12-1F", "ESES12-2", "ESES12-3", "ESES12-4", "ESES12-5M", "ESES12-6F", "ESES13-1F", "ESES13-2F", "ESES13-3F", "ESES13-4F", "ESES13-6", "ESES14-10F", "ESES14-1F", "ESES14-2M", "ESES14-3M", "ESES14-4F", "ESES14-5M", "ESES14-6M", "ESES14-7F", "ESES14-8M", "ESES14-9F", "ESES15-1F", "ESES15-2F", "ESES15-3F", "ESES15-4M", "ESES15-5M", "ESES15-6M", "ESES15-7M", "ESES16-1F", "ESES16-2M", "ESES16-3F", "ESES16-4F", "ESES16-5M", "ESES16-6M", "ESES16-7M", "ESES16-8F", "ESES17-1M", "ESES17-2F", "ESES17-3M", "ESES17-4F", "ESES17-5F", "ESES17-6F", "ESES17-7F", "ESES18-1F", "ESES18-2F", "ESES18-3F", "ESES18-4F", "ESES18-5M", "ESES18-6M", "ESES18-7M", "ESES19-1", "ESES19-2", "ESES19-3", "ESES19-4", "ESES19-5", "ESES19-6", "ESES19-7", "ESES19-8", "ESES19-9", "ESES2-1M", "ESES2-2F", "ESES2-3M", "ESES2-4M", "ESES2-5M", "ESES20-1", "ESES20-2", "ESES20-3", "ESES20-4", "ESES20-5", "ESES20-6", "ESES20-7", "ESES21-1", "ESES21-2", "ESES21-3", "ESES21-4", "ESES21-5", "ESES21-6", "ESES22-1", "ESES22-10", "ESES22-2", "ESES22-3", "ESES22-4", "ESES22-5", "ESES22-6", "ESES22-7", "ESES22-8", "ESES22-9", "ESES3-1M", "ESES3-2F", "ESES3-3F", "ESES3-4M", "ESES4-1F", "ESES4-2F", "ESES4-3M", "ESES5-1M", "ESES5-2M", "ESES5-3M", "ESES5-4M", "ESES5-5F", "ESES5b-1M", "ESES5b-2M", "ESES5b-3", "ESES6-1M", "ESES6-2M", "ESES6-3M", "ESES6-4F", "ESES6-5F", "ESES6-6M", "ESES6-7F", "ESES7-1F", "ESES7-2F", "ESES7-3F", "ESES7-4F", "ESES8-1F", "ESES8-2F", "ESES9-1F", "ESES9-2M", "ESES9-3F", "ESES9-4M", "ESES9-5M", "SSSS0-1M");names(covb)<-Names;covb_backup<-covb
covb<-covb_backup

sexes<-c("F", "F", "F", "M", "M", "F", "F", "M", "M", "F", "M", "M", "M", "M", "M", "M", "F", "M", "F", "F", "F", "M", "NA", "NA", "NA", "NA", "F", "NA", "NA", "NA", "M", "F", "F", "F", "F", "F", "NA", "F", "F", "M", "M", "F", "M", "M", "F", "M", "F", "F", "F", "F", "M", "M", "M", "M", "F", "M", "F", "F", "M", "M", "M", "F", "M", "F", "M", "F", "F", "F", "F", "F", "F", "F", "F", "M", "M", "M", "NA", "NA", "NA", "NA", "NA", "NA", "NA", "NA", "NA", "M", "F", "M", "M", "M", "NA", "NA", "NA", "NA", "NA", "NA", "NA", "NA", "NA", "NA", "NA", "NA", "NA", "NA", "NA", "NA", "NA", "NA", "NA", "NA", "NA", "NA", "NA", "M", "F", "F", "M", "F", "F", "M", "M", "M", "M", "M", "F", "M", "M", "NA", "M", "M", "M", "F", "F", "M", "F", "F", "F", "F", "F", "F", "F", "F", "M", "F", "M", "M", "M") #Order should be good now 16-09-2018 
chr<-covb$chr
if(FALSE){#TRUE makes it just the P and F1s to do sexes for, FALSE makes it everyone
	cov<-covb[,c(3:11,149)];sexes<-sexes[c(1:9,147)]#for only P and F1
}else{cov<-covb[,3:149]} #for all individuals


seq_effort=colSums(cov);head(seq_effort)
cov2<-cov
for(i in 1:ncol(cov)){
	cov2[,i]<-cov[,i]/seq_effort[i]*1000000	
} 
cov3=NULL
Male<-rowMeans(cov2[,sexes=="M"])
Female<-rowMeans(cov2[,sexes=="F"])
cov3<-cbind.data.frame(chr, Male, Female);#head(cov3)
	plot(x=cov3$Female, y=cov3$Male, cex=.1, pch=19, xlim=c(0,100), ylim=c(0,100), bty="L", xlab="Standardized coverage in females", ylab="Standardized coverage in males")


if(FALSE){

	plot(x=cov3$Female, y=cov3$Male, cex=.1, pch=19, bty="L", xlab="Standardized coverage in females", ylab="Standardized coverage in males")
	plot(x=cov3$Female, y=cov3$Male, cex=.1, pch=19, xlim=c(0,100), ylim=c(0,100), bty="L", xlab="Standardized coverage in females", ylab="Standardized coverage in males")
	abline(a=-1.5, b=3/4, col="red")
	segments(-20, 2, .5, 2, col="red")
	segments(100, 800, .5,2, col="red")


	#check the assignments:
#Y:	
	points(cov3$Female[cov3$chr %in% Y[1:100]], cov3$Male[cov3$chr %in% Y[1:100]],col="red", pch=19)
#X:
	points(cov3$Female[cov3$chr %in% X[1:100]], cov3$Male[cov3$chr %in% X[1:100]],col="blue", pch=19)
#A:
	points(cov3$Female[cov3$chr %in% A[1:100]], cov3$Male[cov3$chr %in% A[1:100]],col="green", pch=19)
}

#find the cut-off points. 
U<-cov3$chr[cov3$Male+cov3$Female<3]; length(U)
Y<-cov3$chr[cov3$Female<2 & cov3$Male>4 & ! (cov3$chr %in% U)]; length(Y)
X<-cov3$chr[(3*cov3$Female)-(4*cov3$Male)>(1*4) & ! (cov3$chr %in% U)]; length(X)
A<-cov3$chr[! ((cov3$chr %in% Y) | (cov3$chr %in% X) | (cov3$chr %in% U))]; length(A)
length(U)+length(A)+length(Y)+length(X)==nrow(cov3) #NEEDS to be TRUE!!

covY<-cov2[chr%in%Y,] #
plot(colMeans(covY), col=as.factor(sexes), pch=paste(c(1:9,0)));legend(x=0,y=3, legend=c("Male", "Female", "unkown"), fill=c(2,1,3))


#######################################################
#since there were a couple mis-sexed individuals, it's now time to re-call sexes, re-plot the figure, and re-call linkage

new_sexes<-rep("F", length(sexes)); new_sexes[colMeans(covY)>3]<-"M"
names(new_sexes)<-colnames(covY)
plot(colMeans(covY), col=as.factor(new_sexes), pch=paste(0:9));legend(x=0,y=3, legend=c("Male", "Female", "unkown"), fill=c(2,1,3))

cov4=NULL
Male<-rowMeans(cov2[,new_sexes=="M"])
Female<-rowMeans(cov2[,new_sexes=="F"])
cov4<-cbind.data.frame(chr, Male, Female);#head(cov4)
	plot(x=cov4$Female, y=cov4$Male, cex=.1, pch=19, xlim=c(0,100), ylim=c(0,100), bty="L", xlab="Standardized coverage in females", ylab="Standardized coverage in males")


Y<-cov4$chr[cov4$Male-5*cov4$Female>.2]; length(Y)
U<-cov4$chr[cov4$Male+cov4$Female<1 & ! (cov4$chr %in% Y)]; length(U)
X<-cov4$chr[cov4$Male-(7/10)*cov4$Female<(-.05) & ! (cov4$chr %in% U)]; length(X)
A<-cov4$chr[! ((cov4$chr %in% Y) | (cov4$chr %in% X) | (cov4$chr %in% U))]; length(A)
length(U)+length(A)+length(Y)+length(X)==nrow(cov3) #NEEDS to be TRUE!!

if(TRUE){
	#plot(x=cov3$Female, y=cov3$Male, cex=.1, pch=19, bty="L", xlab="Standardized coverage in females", ylab="Standardized coverage in males")
	plot(x=cov4$Female, y=cov4$Male, cex=.1, pch=19, xlim=c(0,10), ylim=c(0,10), bty="L", xlab="Standardized coverage in females", ylab="Standardized coverage in males")
	abline(a=-0.05, b=7/10, col="red") # X-A boundary, less = X, more=A
	abline(a=0.2, b=5, col="blue") # Y-A boundary, less=A, more=Y
	abline(a=1, b=-1, col="gray") # U-A/X boundary, less = U, more=A/X
	#check the assignments:
	points(cov4$Female[cov4$chr %in% U[1:100]], cov4$Male[cov4$chr %in% U[1:100]],col="gray", pch=19) #U
	points(cov4$Female[cov4$chr %in% Y[1:100]], cov4$Male[cov4$chr %in% Y[1:100]],col="red", pch=19) #Y
	points(cov4$Female[cov4$chr %in% X[1:100]], cov4$Male[cov4$chr %in% X[1:100]],col="blue", pch=19) #X
	points(cov4$Female[cov4$chr %in% A[1:100]], cov4$Male[cov4$chr %in% A[1:100]],col="green", pch=19) #A
}


#Xscaf<-read.table("NW_018657747.1_rads.txt", header=F)$V1
#cov5<-cov4[cov4$chr%in%Xscaf,]
#points(cov5$Female, cov5$Male, col="red", pch=19)

#######################################################
if(FALSE){ #didn't end up using this - it tries to look at the averages across each scaffold, ended up not helping much so I skipped it in the final version. 
library(data.table)
#Now refine the predicitons based on scaffold-wide averages
sam<-read.table("catalog_GCF.partsam", header=F, stringsAsFactors=T);names(sam)<-c("rad", "flag", "scaf", "pos", "qual", "cigar");head(sam)
cov5<-cov4
cov5$Female<-ifelse(cov5$Female==0,0.0000000001, cov5$Female)
cov5$ratio=cov5$Male/cov5$Female;head(cov5)
sam2<-merge(x=sam, y=cov5, by.x="rad", by.y="chr", all.x=F, all.y=F)

sam_list<-split(sam2, f=sam2$scaf)

myFun<-function(df){
	
	medianRatio<-rep(median(df$ratio, na.rm=T), nrow(df))
	meanMale<-rep(mean(df$Male, na.rm=T), nrow(df))
	meanFemale<-rep(mean(df$Female, na.rm=T), nrow(df))
	df<-cbind(df,meanMale, meanFemale, medianRatio)
}

sam_list<-lapply(sam_list, FUN=myFun)
head(sam_list)

cov6<-rbindlist(sam_list[1:100])


#points(sam3$meanFemale[which(sam3$rad%in%Xscaf)], sam3$meanMale[which(sam3$rad%in%Xscaf)], col="red", pch=19)

Y<-cov6$rad[cov6$meanMale-5*cov6$meanFemale>.2]; length(Y)
U<-cov6$rad[cov6$meanMale+cov6$meanFemale<1 & ! (cov6$rad %in% Y)]; length(U)
X<-cov6$rad[cov6$meanMale-(7/10)*cov6$meanFemale<(-.05) & ! (cov6$rad %in% U)]; length(X)
A<-cov6$rad[! ((cov6$rad %in% Y) | (cov6$rad %in% X) | (cov6$rad %in% U))]; length(A)
length(U)+length(A)+length(Y)+length(X)==nrow(cov6) #NEEDS to be TRUE!!
}


#write out the data!
if(TRUE){
	write.table(x=A, file="Autosomal.txt", quote=F, row.names=F, col.names=F)
	write.table(x=X, file="Xlinked.txt", quote=F, row.names=F, col.names=F)
	write.table(x=Y, file="Ylinked.txt", quote=F, row.names=F, col.names=F)
	write.table(x=U, file="Unknown.txt", quote=F, row.names=F, col.names=F)
}


#make the figures!
if(TRUE){
png(file="~/Dropbox/Post_Doc/Manuscripts/Meriones_Map/figures/Figure_2_Sex_Chr_Annotation/Figure_2a_Sex_Chr_Annotation.png", width=7, height=7, units="in", res=500)
plot(x=cov4$Female, y=cov4$Male, cex=.3, pch=19, bty="L", xlim=c(0,600), ylim=c(0,600), xlab="Standardized coverage in females", ylab="Standardized coverage n males", type="n", main="GBStag coverage - not scaffolds")
points(x=cov4$Female[cov4$chr %in% X], y=cov4$Male[cov4$chr %in% X], cex=.1, pch=20,col="red")
points(x=cov4$Female[cov4$chr %in% Y], y=cov4$Male[cov4$chr %in% Y], cex=.1, pch=20,col="blue")
points(x=cov4$Female[cov4$chr %in% A], y=cov4$Male[cov4$chr %in% A], cex=.1, pch=20,col="black")
points(x=cov4$Female[cov4$chr %in% U], y=cov4$Male[cov4$chr %in% U], cex=.1, pch=20,col="grey")
legend(x=450, y=200, legend=c("Autosomal", "X-linked", "Y-linked", "Unknown"), fill=c("black", "red", "blue", "grey"))
dev.off()


#zoom in with histograms:
png(file="~/Dropbox/Post_Doc/Manuscripts/Meriones_Map/figures/Figure_2_Sex_Chr_Annotation/Figure_2b_Sex_Chr_Annotation.png", width=7, height=7, units="in", res=500)
xhist<-hist(cov4$Female,breaks=c(0:12973), plot=F)
yhist<-hist(cov4$Male, breaks=c(0: 12914), plot=F)
nf<-layout(matrix(c(2,0,1,3),2,2,byrow=T), c(8,1), c(1,8), TRUE) #specifies the layout of the plot area
par(mar=c(5,5,.5,.5))
plot(x=cov4$Female, y=cov4$Male, pch=19, xlim=c(0,80), ylim=c(0,80), bty="L", xlab="Average Standardized coverage in females", ylab="Average standardized coverage in males", type="n", xaxs="r", yaxs="r", bty="n")
if(T){
	points(x=cov4$Female[cov4$chr %in% A], y=cov4$Male[cov4$chr %in% A], cex=.5, pch=3, col="black")#pch=20, col="black")
	points(x=cov4$Female[cov4$chr %in% X], y=cov4$Male[cov4$chr %in% X], cex=.7, pch=16, col="grey30")#pch=20, col="red")
	points(x=cov4$Female[cov4$chr %in% Y], y=cov4$Male[cov4$chr %in% Y], cex=.7, pch=17, col="grey 60")#pch=20, col="blue")
	points(x=cov4$Female[cov4$chr %in% U], y=cov4$Male[cov4$chr %in% U], cex=.7, pch=18, col="grey")#pch=20, col="grey")
}
legend(x=10, y=60, legend=c("Autosomal", "X-linked", "Y-linked", "Unknown"), pch=c(3,16,17,18), col=c("black", "grey30", "grey60", "grey"))#, fill=c("black", "red", "blue", "grey"))
#abline(v=c(0,100), h=c(0,100))
par(mar=c(0,4.3,1,0))
barplot(1*xhist$counts[1:100], axes=F, xlim=c(1,100), space=0, horiz=F)

par(mar=c(4.3,0,0,1))
barplot(1*yhist$counts[1:100], axes=F, ylim=c(1,100), space=0, horiz=T)

dev.off()#abline(a=-5, b=24/30, col="red")
#segments(-20, 10, 1, 10, col="red")
#segments(2, 800, 1,10, col="red")

#Show the cutoffs - super zoom in
png(file="~/Dropbox/Post_Doc/Manuscripts/Meriones_Map/figures/Figure_2_Sex_Chr_Annotation/Figure_2c_Sex_Chr_Annotation.png", width=7, height=7, units="in", res=500)
plot(x=cov4$Female, y=cov4$Male, pch=19, xlim=c(0,5), ylim=c(0,5), bty="L", xlab="Average Standardized coverage in females", ylab="Average standardized coverage in males", type="n", xaxs="r", yaxs="r", bty="n")
if(T){
	points(x=cov4$Female[cov4$chr %in% A], y=cov4$Male[cov4$chr %in% A], cex=.05, pch=20,col="black")
	points(x=cov4$Female[cov4$chr %in% X], y=cov4$Male[cov4$chr %in% X], cex=.05, pch=20,col="red")
	points(x=cov4$Female[cov4$chr %in% Y], y=cov4$Male[cov4$chr %in% Y], cex=.05, pch=20,col="blue")
	points(x=cov4$Female[cov4$chr %in% U], y=cov4$Male[cov4$chr %in% U], cex=.05, pch=20,col="grey")
}
abline(a=-0.05, b=7/10, col="red", cex=.5, lty=2) # X-A boundary, less = X, more=A
abline(a=0.2, b=5, col="blue", cex=.5, lty=2) # Y-A boundary, less=A, more=Y
abline(a=1, b=-1, col="gray", cex=.5, lty=2) # U-A/X boundary, less = U, more=A/X
text("Autosomal", cex=1, x=2, y=4.5)
text("Y-linked", cex=1, x=.5, y=4.9, col="blue")
text("X-linked", cex=1, x=4, y=0.8, col="red")
text("Unknown", cex=1, x=.55, y=0, col="gray50")
dev.off()


}
```
