## Supplemental_File_5_Codebase for "A high-density genetic map and molecular sex-typing assay for gerbils": evaluate_and_format_distance_matrix.R

```
#!/usr/bin/env Rscript
#29-05-2018

#based on the one for the phodopus project, this script goes through and places the heterozygous markers on the map for hte meriones project
#this script should go through and format the locations of the capture markers into a 5-column file:
#marker_name	chr	cM	
#to be run on a computer with dropbox

setwd("/Volumes/LaCie/Projects/Meriones_Map/stacks_out")
dis1<-read.table("het_marker_dist_matrix.txt", header=T, stringsAsFactors=F);dim(dis1)#for distance matrix: formed by command: make_rqtl_input_file_for_Capture_2.py -v ES_Var.recode.vcf -m ../stacks_out/Meriones_Map.csv -o het_marker_dist_matrix.txt -c 70 
dis1SA<-read.table("het_marker_dist_matrix.SWITCHED_ALLELES.txt", header=T, stringsAsFactors=F);dim(dis1)
dis2<-read.table("hom_marker_dist_matrix.txt", header=T, stringsAsFactors=F);dim(dis2)#hom_marker_dist_matrix.txt
dis2SA<-read.table("hom_marker_dist_matrix.SWITCHED_ALLELES.txt", header=T, stringsAsFactors=F);dim(dis2SA)#hom_marker_dist_matrix.txt

dis<-cbind(dis1,dis2[,3:ncol(dis2)]);dim(dis)

#############
#############

palette(c("black",   "red" ,    "green3" , "blue"  ,  "cyan"   , "magenta" ,"yellow"  ))

#this bit makes all the plot files for the gif
if(FALSE){
	counter=0
	for (i in 3:ncol(dis)){
		cat("Plotting file", i, "\r")
		if (i < 10) {name = paste('000',i,'_',colnames(dis)[i],'.png',sep='')} 
		if (i < 100 && i >= 10) {name = paste('00',i,'_',colnames(dis)[i],'.png', sep='')} 
		if (i >= 100) {name = paste('0', i,'_',colnames(dis)[i],'.png', sep='')}
		if (i >= 1000) {name = paste( i,'_',colnames(dis)[i],'.png', sep='')}
		if (i >= 1000) {name = paste(i,'_',colnames(dis)[i],'.png', sep='')}
		png(paste0("../het_markers/Position_plots/unplaced/", name))
		themean=mean(dis[,i]);themean
		thesd=sd(dis[,i]);thesd
		themin=min(dis[,i]);themin
		if (themin>.1){
			counter=counter+1
			plot(y=dis[,i], x=1:nrow(dis), pch=19, cex=.1, xlab="genomic location", ylab="distance", main=colnames(dis)[i], ylim=c(0,1), type="n", bty="L")
			rect(xleft=0,xright=44,ybottom=0,ytop=100,col="grey90", border=NA)
			rect(xleft=100,xright=151,ybottom=0,ytop=100,col="grey90", border=NA)
			rect(xleft=179,xright=207,ybottom=0,ytop=100,col="grey90", border=NA)
			rect(xleft=229,xright=255,ybottom=0,ytop=100,col="grey90", border=NA)
			rect(xleft=263,xright=305,ybottom=0,ytop=100,col="grey90", border=NA)
			rect(xleft=339,xright=360,ybottom=0,ytop=100,col="grey90", border=NA)
			rect(xleft=373,xright=386,ybottom=0,ytop=100,col="grey90", border=NA)
			rect(xleft=405,xright=417,ybottom=0,ytop=100,col="grey90", border=NA)
			rect(xleft=430,xright=444,ybottom=0,ytop=100,col="grey90", border=NA)
			rect(xleft=457,xright=468,ybottom=0,ytop=100,col="grey90", border=NA)
			rect(xleft=475,xright=482,ybottom=0,ytop=100,col="grey90", border=NA)
			points(y=dis[,i], x=1:nrow(dis), pch=19, cex=.1, type="l", xlab="genomic location", ylab="distance", main=colnames(dis)[i], ylim=c(0,1));
			points(y=dis[,i], x=1:nrow(dis), pch=19, cex=.3, type="p", xlab="genomic location", ylab="distance", main=colnames(dis)[i], ylim=c(0,1), col=as.factor(dis$Chr));
			abline(h=themean, col="gray", lty=2)
			abline(h=themean-(4*thesd), col="orange", lty=3)
			abline(h=.1, col="red", lty=2)
		}		
	dev.off()
	}
}


####################################################################################
#now make the list of markers that fit somewhere with min<.1:
out<-data.frame(matrix(rep(NA,(100000*4)),ncol=4));colnames(out)=c("RADID", "Chr", "cM", "SA");

#for hets:
j=0
for(i in 3:ncol(dis1)){
	themin=min(dis1[,i]);themin
	if (themin<.1){
		#cat(i, "	")
		j=j+1
		idx=which(dis1[,i]==min(dis1[,i]))[1];idx
		out$RADID[j]<-colnames(dis1)[i]
		out$Chr[j]<-dis1$Chr[idx]
		out$cM[j]<-dis1$cM[idx]	
		out$SA[j]<-"NO"
	}else {
		themin=min(dis1SA[,i]);themin
		if(themin<.1){
			j=j+1
			idx=which(dis1SA[,i]==min(dis1SA[,i]))[1];idx
			out$RADID[j]<-colnames(dis1SA)[i]
			out$Chr[j]<-dis1SA$Chr[idx]
			out$cM[j]<-dis1SA$cM[idx]
			out$SA[j]<-"YES"
	
		}
	}
}
out=out[complete.cases(out),]
out=out[order(out[,2],out[,3],out[,1]),]
outhet<-out;head(outhet)

#for homs:
out<-data.frame(matrix(rep(NA,(100000*4)),ncol=4));colnames(out)=c("RADID", "Chr", "cM", "SA");head(out)
j=0
for(i in 3:ncol(dis2)){
	themin=min(dis2[,i]);themin
	if (themin<.1){
		j=j+1
		idx=which(dis2[,i]==min(dis2[,i]))[1];idx
		out$RADID[j]<-colnames(dis2)[i]
		out$Chr[j]<-dis2$Chr[idx]
		out$cM[j]<-dis2$cM[idx]
		out$SA[j]<-"NO"
	
	}else{
		themin=min(dis2SA[,i]);themin
		if (themin<.1){
			j=j+1
			idx=which(dis2SA[,i]==min(dis2SA[,i]))[1];idx
			out$RADID[j]<-colnames(dis2SA)[i]
			out$Chr[j]<-dis2SA$Chr[idx]
			out$cM[j]<-dis2SA$cM[idx]
			out$SA[j]<-"YES"
	
		}	
	}
}
out=out[complete.cases(out),]
out=out[order(out[,2],out[,3],out[,1]),]
outhom<-out;dim(outhom);#head(outhom)

dim(outhet)
dim(outhom)


write.table(x=outhet, file="../stacks_out/het_Markers_placed.txt", quote=F, sep="	", row.names=F, col.names=F)
write.table(x=outhom, file="../stacks_out/hom_Markers_placed.txt", quote=F, sep="	", row.names=F, col.names=F)

write(date(), file="evaluate_and_formate_distance_matrix.R.finished")


####################################################################################
```
