## Supplemental_File_5_Codebase for "A high-density genetic map and molecular sex-typing assay for gerbils": extract_fasta_region.py

```
#!/usr/bin/env Python3


import sys
import math
import argparse
from Bio.SeqIO.FastaIO import SimpleFastaParser


parser = argparse.ArgumentParser(description="written by TDB\n10-01-2018\nThis script takes a fasta and extracts the desired bits from it. Input can be a (1) single entry with -chr, (2) a single entry, a start, and a stop, with --chr -s -e, (3) a list of entries with --chrList, or (4) a list of entries, starts, and stops in three tab-separated columns also with --chrList. It makes a new fasta of just the bases thus described.")
parser.add_argument("-f", "--Fasta", type=str, required=True, help="A fasta file. Default: %(default)s", default="", metavar="*.fasta")
parser.add_argument("-chr", "--chromosome", type=str, required=False, help="The chromosome to be extracted")
parser.add_argument("-s", "--start", type=str, required=False, help="A start base. Indexed such that the first base in the sequence is 1 (not 0). Default:  %(default)s", metavar="int", default=1)
parser.add_argument("-e", "--end", type=str, required=False, help="An end base. Default: sequence length", metavar="int", default=100000000000000000)
parser.add_argument("-chrList", "--chromosome_List", type=str, required=False, help="A tab-sparated text file that has a list of chr names to be extracted. Optionally has a column of starts and a column of stops for each chromosome. The file may also have an optional second or fourth column of strand ('+' or '-'). 1 column = chr, 2 columns = chr strand, 3 columns = chr start end, 4 columns = chr start end strand. Chromosomes may appear multiply, but only one end base can go with each start base.")
parser.add_argument("-o", "--out", type=str, required=False, help="A output file name. Missing goes to stdout", metavar="*.fasta")
parser.add_argument("-st", "--strand", type=str, required=False, help="The strand from the fasta that the bases should be pulled from: + or -. Should only be used with '-chr', not '-chrList'. Default: %(default)s", default='+')
parser.add_argument("-fmt", "--outfmt", type=str, required=False, help="The number of bases per line the new fasta should have. Default: %(default)s", default="100",)


args = parser.parse_args()
s=int(args.start)
e=int(args.end)
start=min(s,e)
end=max(s,e)
fmt=int(args.outfmt)
chr=args.chromosome
out=args.out


if out is None:
	OUT=sys.stdout
else:
	OUT=open(out, 'w')

def revcom(str):
	rcd = {"A":"T", "G":"C", "C":"G", "T":"A", "a":"t", "g":"c", "c":"g", "t":"a", "N":"N", "n":"n"}
	com = "".join([rcd[b] for b in str])
	rcom = com[::-1]
	return(rcom)

def parse_chrList(listfile):
	with open(listfile, 'r') as LIST:
		for item in LIST:
			item=item.strip("\n")
			item=item.rstrip()
			item=item.lstrip(">")
			item=str(item)
			#print(item)
			bits=item.split("	")
			#print(len(bits), file=sys.stderr)
			if len(bits)==2:
				chr=bits[0]
				strand=bits[1]
				if chr not in chr_list:
					chr_list[chr]["region"]={1:{1000000000000000000000000000000000000:strand}}
					#chr_list[chr]["strand"]=strand
			elif len(bits)>=3:
				chr=bits[0]
				start_init=int(bits[1])
				stop_init=int(bits[2])
				start = min(start_init, stop_init)
				stop = max(start_init, stop_init)
				if chr not in chr_list:
					chr_list[chr]={"region":{}}
				chr_list[chr]["region"][start] = {stop : "+"}
				
				if len(bits)==4:
					strand=bits[3]
					chr_list[chr]["region"][start][stop]=strand
				else:
					chr_list[chr]["region"][start][stop]="+"	
			elif len(bits)==1:	
				chr=bits[0]	
				if chr not in chr_list:
					chr_list[chr]={}
				chr_list[chr]["region"]={1:{1000000000000000000000000000000000000:"+"}}
				#chr_list[chr]["strand"]="+"
			else:
				print("\nERROR: The chromosome_List is formatted incorrectly - it needs three tab separated columns of chr_name \t start \t stop. Start and stop should be base-1 indexed.", file=sys.stderr, end="\n\n")
	
ROILength=end-start
nlines=math.ceil(ROILength/fmt)

chr_list={}
if args.chromosome_List is not None:
	parse_chrList(args.chromosome_List)
elif args.chromosome is None:	
	print("\nERROR: Needs at least either a chromosome (-chr) or a chromosome list (-chrList)", end="\n\n", file=sys.stderr)
	exit()
else:
	if args.strand:
		strand = args.strand
	else:
		strand = "+"
		
	chr_list[args.chromosome]={}
	chr_list[args.chromosome]["region"]={start:{end:strand}}
	#chr_list[args.chromosome]["strand"]=args.strand
#print(chr_list, file=sys.stderr)

#print(chr_list)

with open(args.Fasta, "r") as FASTAH:
	for record in SimpleFastaParser(FASTAH): 
		name=record[0].split()[0]
		print("Scanning fasta: ", name, "len =", len(record[1]), file=sys.stderr, end="\r")
		if name in chr_list:
			#print(name, "is in chr_list:", chr_list, file=sys.stderr)
			for start in chr_list[name]["region"]:
				end = [*chr_list[name]["region"][start]][0]
				end1 = end
				if end > len(record[1]):
					end = len(record[1])
				seq=record[1][int(start)-1:end] #the sequence - from desired start to desired stop. 
				if chr_list[name]["region"][start][end1]=="-":
					seq=revcom(seq)
					start,end=end,start
				print(">", name, "	EXTRACTED REGION: bases	", start, ":", end, sep="", file=OUT)
				lines_to_print=int(min(nlines, (len(seq)/fmt)))
				for i in range(lines_to_print+1):
					low=i*fmt
					high=((i+1)*fmt)-1
					print(seq[low:high+1], file=OUT)		
if out is not None:
	OUT.close()		
print("																	", file=sys.stderr, end="\r")	
	
```
