## Supplemental_File_5_Codebase for "A high-density genetic map and molecular sex-typing assay for gerbils": filter_SNPs.py

```
#!/usr/bin/env Python3

import sys
import argparse

parser = argparse.ArgumentParser(description="written by TDB\n23-11-2017\nSpecifically for the Meriones_Map project.\nFilters out sites that are fixed in one parent alternativly from the other. Also pull out ones fixed in one, but heterozygous in the other.")                                              
parser.add_argument("-v", "--VCF", type=str, required=False, help="A vcf file. Default: %(default)s", default="catalog.calls.p.snps.vcf", metavar="*.vcf")
parser.add_argument("-o1", "--OUT1", type=str, required=False, help="A vcf file. Default: %(default)s", default="ES_Fixed.vcf", metavar="*.vcf")
parser.add_argument("-o2", "--OUT2", type=str, required=False, help="A vcf file. Default: %(default)s", default="ES_Var.vcf", metavar="*.vcf")
args = parser.parse_args()
VCF = args.VCF
OUT1 = args.OUT1
OUT2 = args.OUT2


OUT1=open(OUT1, 'w')
OUT2=open(OUT2, 'w')
fixed=0
var=0
badlines=0
oddlines=0


def parse_line(str):
	fields=str.split()
	#print(fields)
	E=fields[9] #The EEEE parent - cut -f 10 <-- python base-0
	S=fields[155] #The SSSS parent - cut -f 156 <--python is base 0
	out="odd"
	if E=="./." or S=="./.":
		out="Bad"
	else:	
		Egeno=E.split(":")[0]
		Sgeno=S.split(":")[0]
		
		if Sgeno==Egeno:
			out="Bad"
		else:
			if Sgeno == "0/1" or Egeno == "0/1":
				out="Var"	
			else:
				out="Fixed"
		#print(Egeno,Sgeno, out)
	return(out)

with open(VCF, 'r') as VCF:
	for line in VCF:
		if line.startswith("#"):
			print(line, end="", file=OUT1)
			print(line, end="", file=OUT2)
		else:
			flag=parse_line(line)	#flag will come out as Fixed, Var, or Bad
			if flag=="Fixed":
				print(line, end="", file=OUT1)
				fixed+=1
			elif flag=="Var":
				print(line, end="", file=OUT2)
				var+=1
			elif flag=="Bad":
				badlines+=1
			else:
				oddlines+=1
				print("for some reason the parser falied - flag is not Fixed, Var or Bad. Instead it is:", flag, file=sys.stderr)	
print("Alternatively Fixed:	", fixed, sep="	", file=sys.stderr)
print("Variable in one parent:	", var, sep="	", file=sys.stderr)
print("No diversity in parents:", badlines, sep="	", file=sys.stderr)
if oddlines:
	print("Odd lines - something else:", oddlines, sep="	", file=sys.stderr)
				
				
```
