## Supplemental_File_5_Codebase for "A high-density genetic map and molecular sex-typing assay for gerbils": get_details_of_mapping.py

```
#!/usr/bin/env Python3


import sys
import re
import argparse
expl='''
get_details_of_mapping.py
created by TDB 14-08-2014

Takes in the map, an alignment file, and a gff and outputs some stats - how many genes on the map and how many bases on the map. 
Also outputs a list of the genes on the map 
'''
parser=argparse.ArgumentParser(description=expl)
parser.add_argument("--gff", "-g", type=str, required=True, help="The GFF of the meriones genome.")
parser.add_argument("--sam", "-s", type=str, required=True, help="The .sam alignment of the catalog to the GCF genome with a sam header")
parser.add_argument("--map", "-m", type=str, required=True, help="The Genetic map of GBS genotypes from gerbils - Meriones_Map_AllMarkers_final.csv perhaps")
parser.add_argument("--out", "-o", type=str, required=False, help="Outfile name. This will be where the list of genes goes. Omit for stdout")
#parser.add_argument("--CSVR", default=None, action='store_const', const=True, required=False, help="Forces script to make a csvr (lacking a header). Used to add genotypes to the map. Will be named after value declared by --out with .csvr as the stem.")

args=parser.parse_args()


if args.out:
	OUT=open(args.out, "w")
else:
	OUT=sys.stdout


def parse_sam(sam):
	with open(sam, 'r') as SAM:
		for line in SAM:
			if not line.startswith("@SQ"):
				return("")
			line = line.strip("\n")
			bits = re.search("^@SQ\s+SN:(.+)\s+LN:(.+)$",line)
			if bits.group(1) not in contig_dict:
				contig_dict[bits.group(1)] = {}
			contig_dict[bits.group(1)].update({"Length":int(bits.group(2))})	

def parse_map_name(name, what = "all"):
	if what == "all":	
			return(name)		
	bits = name.split("_")
	#print(bits)
	if len(bits) == 6:
		contig = "_".join(bits[0:2])
		contig_pos = bits[2]
		rad = bits[3]
		rad_pos = bits[4]
		#print(contig)
		if what == "contig":
			return(contig)
		elif what == "rad":
			return(rad)	
	elif len(bits) == 2:
		rad = bits[0]
		rad_pos = bits[1]
		if what == "contig":
			return("None")
		elif what == "rad":
			return(rad)

def parse_gff_line(line):
	#print(line, end="")
	bits=line.split()
	contig=bits[0]
	if contig not in contig_dict:
		contig_dict[contig]={"Length":0}
	info=bits[8]
	infobits=info.split(";")
	gene=infobits[0].split("=")[1]
	if "Genes" not in contig_dict[contig]:
		contig_dict[contig].update({"Genes":{}})
		contig_dict[contig].update({"GeneCount":0})
	for i in infobits[1:]:
		I=i.split("=")
		if gene not in contig_dict[contig]["Genes"]:
			contig_dict[contig]["Genes"].update({gene:{}})
			contig_dict[contig]["GeneCount"]+=1	
		contig_dict[contig]["Genes"][gene].update({I[0]:I[1]})


def parse_gff(gff):
	with open(gff,'r') as GFF:
		for line in GFF:
			if not line.startswith("#") and re.search("	gene	", line):
				parse_gff_line(line)

			
def parse_map(map):
	linecounter=0
	with open(map, 'r') as MAP:
		for line in MAP:
			linecounter+=1
			#print(line.strip("\n"))
			if linecounter>7:
				name = line.split(",")[0]
				contig = parse_map_name(name, what = "contig")	
				if contig not in map_dict:
					map_dict[contig]=0
				map_dict[contig]+=1	
				
				


def get_stats():
	thelen = 0
	totallen = 0
	contigs1MB = 0
	contigs01MB = 0

	geneCount = 0
	totalgenes = 0
	
	for contig in map_dict:
		if map_dict[contig] = 1:
			thelen += contig_dict[contig]["Length"]
			if "GeneCount" in contig_dict[contig]: # if the contig has genes in it.
				geneCount += contig_dict[contig]["GeneCount"]
				for gene in contig_dict[contig]["Genes"]: 
					print(contig_dict[contig]["Genes"][gene]["Name"], file=OUT)		
		else:
			print(contig, "is in the genome multiple times...", file=sys.stderr)
	for i in contig_dict:
		totallen += contig_dict[i]["Length"]
		if "GeneCount" in contig_dict[i]:
			totalgenes += contig_dict[i]["GeneCount"]
		if i in map_dict and contig_dict[i]["Length"]>=1000000:
			contigs1MB += 1
		if i in map_dict and contig_dict[i]["Length"]>=100000:
			contigs01MB += 1
	
	lenpercent = round(thelen*100/totallen, 2)	
	genepercent = round(geneCount*100/totalgenes,2)	
	scafpercent = round(len(map_dict)*100/len(contig_dict),2)	
	contigs1MBpercent = round(contigs1MB*100/len(map_dict), 2)		
	contigs01MBpercent = round(contigs01MB*100/len(map_dict), 2)		

	print("\nTotal number of bases associated with map:	", thelen, " = ", lenpercent, "% of genomic bases", sep="", file=sys.stderr)	
	print("Total number of genes associated with map:	", geneCount, " = ", genepercent, "% of annotated genes", sep="", file=sys.stderr)	
	print("Number of scaffolds contigs associated with map:	", len(map_dict), " = ", scafpercent, "% of scaffolds", sep="",file=sys.stderr)		
	print("Number of scaffolds contigs > 1.0Mb associated with map:	", contigs1MB, " = ",contigs1MBpercent, "% of placed scaffolds", sep="",file=sys.stderr, end="\n")		
	print("Number of scaffolds contigs > 0.1Mb associated with map:	", contigs01MB, " = ",contigs01MBpercent, "% of placed scaffolds", sep="",file=sys.stderr, end="\n\n")		

	#print(scaffs)		
			
				
contig_dict = {"None":{"Length":0}} # this is a complex dictionary of dictionaries including length and annotation information:
									#contig_dict  	- "Length" : 123 scaffold length
									#				- "Genes" : many bits of annotation infor includeing gene name and others  			
									#				- "GeneCount" : 12 number of genes
parse_sam(args.sam)			
parse_gff(args.gff)
map_dict = {} # keys are genome contigs, values are the counts that it shows up in the genome. - should be 1...
parse_map(args.map)
get_stats()

#for i in contig_dict:
#	if "Genes" in contig_dict[i]:
#		print(i, contig_dict[i]["Length"], contig_dict[i]["GeneCount"])
		
if args.out:
	OUT.close()
		
		
		
```
