## Supplemental_File_5_Codebase for "A high-density genetic map and molecular sex-typing assay for gerbils": Get_Xlinked_tags.py

```
#!/usr/bin/env	Python3		
import sys
import argparse

#print("this script is deprecated for extract_fasta_region.py", file=sys.stderr)
#exit()

parser = argparse.ArgumentParser(description="written by TDB\n07-05-2018\nSpecifically for the Meriones_Map project.\nPulls out X-linked SNPs from the rqtl input file.")                                              
parser.add_argument("-x", "--Xlinked", type=str, required=True, help="A text file of GBS tags that are x-linked.", metavar="*.txt")
parser.add_argument("-y", "--Ylinked", type=str, required=False, help="A text file of GBS tags that are y-linked.")
parser.add_argument("-r", "--rqtl", type=str, required=True, help="An rqtl file of GBS tags.", metavar="*.rqtl")
parser.add_argument("-r2", "--rqtlVar", type=str, required=True, help="An rqtl file of GBS tags from the Var set: rqtl_input_Var_2.csvr")
parser.add_argument("-o", "--outX", type=str, required=False, help="Output file name, omit for stdout.", metavar="*.txt")
parser.add_argument("-p", "--outY", type=str, required=False, help="Output file name, if omitted, no Y linked tags will be printed.", metavar="*.txt")

args = parser.parse_args()
XFH = args.Xlinked
YFH = args.Ylinked
rqtlFH = args.rqtl
rqtl2FH = args.rqtlVar
Out = args.outX
Out2 = args.outY

if Out2 and YFH:
	OUT2=open(Out,'w')	
if not Out:
	OUT=sys.stdout
else:
	OUT=open(Out, 'w')

Ydict={}
Xdict={}
print("Reading in X linked tags", file=sys.stderr)
with open(XFH, 'r') as X:
	for line in X:
		line=line.strip("\n")
		Xdict[line]=1

if Out2 and YFH:
	print("reading in Y linked tags", file=sys.stderr)
	with open(YFH, 'r') as Y:
		for line in Y:
			line=line.strip("\n")
			Ydict[line]=1	


print("Reading in RQTL file", file=sys.stderr)
with open(rqtlFH, 'r') as RQTL:
	for line in RQTL:
		tagname=line.split(",")[0]
		#print(tagname, file=sys.stderr)
		if tagname.startswith("N"):
			tag=tagname.split("_")[4]
		else:
			tag=tagname.split("_")[0]
		if tag in Xdict:		
			print(tagname, file=OUT)	
		if Out2 and YFH:
			if tag in Ydict:
				print(tagname, file=OUT2)
if rqtl2FH:
	print("Reading in RQTL2 file", file=sys.stderr)
	with open(rqtl2FH, 'r') as RQTL:
        	for line in RQTL:
                	tagname=line.split(",")[0]
                	#print(tagname, file=sys.stderr)
                	if tagname.startswith("N"):
                	        tag=tagname.split("_")[4]
                	else:
                	        tag=tagname.split("_")[0]
                	if tag in Xdict:
                	        print(tagname, file=OUT)
                	if Out2 and YFH:
                	        if tag in Ydict:
                	                print(tagname, file=OUT2)


if Out:
	OUT.close()		
if Out2 and YFH:
	OUT2.close()
```
