## Supplemental_File_5_Codebase for "A high-density genetic map and molecular sex-typing assay for gerbils": make_rqtl_input_file.py

```
#!/usr/bin/env Python3

import argparse
import re
import select
import sys


class TimeoutExpired(Exception):
    pass
def input_with_timeout(prompt, timeout):
    sys.stdout.write(prompt)
    sys.stdout.flush()
    ready, _, _ = select.select([sys.stdin], [],[], timeout)
    if ready:
        return sys.stdin.readline().rstrip('\n') # expect stdin to be line-buffered
    raise TimeoutExpired

    


parser = argparse.ArgumentParser(description="written by TDB\n09-12-2014\nUpdated by TDB on 22-11-2017 for use with any VCF")                                              
parser.add_argument("--VCF", "-v", type=str, required=True, help="VCF file with genotypes. Default: %(default)s", default="gstacks.vcf", metavar="*.vcf")
parser.add_argument("--Pheno", "-p", type=str, required=False, help="If phenotypes need be added use this. It needs to be a tsv formatted file with phenotypes in columns, first column must be the names of the individuals EXACTLY the same as in the VCF OR -t must be used, see below.", metavar="*.txt")
parser.add_argument("--Out", "-o", type=str, required=True, help="Output file name.", metavar="*.csvr")
parser.add_argument("--name_to_Trim", "-t", type=str, required=False, help="If vcf names have an extra bit (i.e. sorted.bam) that you don't want to use, type it here and it will be removed. Perhaps: _R1_clipped_passed-re-filter", metavar="STR")
parser.add_argument("--only_first_SNP", default=None, action='store_const', const=True, required=False, help="Include to only use the first SNP per radtag.")

args = parser.parse_args()

vcf = args.VCF
Pheno = args.Pheno
Out = args.Out
Trim = args.name_to_Trim
OUT=open(Out, 'w')
names=[] # for a list of individual names to be able to call the dict later.


def format_Header(str): #first make the header row for the table: do this from the id's of the vcf file: ID,,ind1,ind2,ind3... followed by a row of sexes: sex,,1,1,1,2,2,1,1...
	fields=str.split()
	print("ID,", sep=",", end="", file=OUT)
	cols=0
	#first line: individual names and make a list of the individuals to use later
	for name in fields[9:]:
		cols+=1
		if Trim: #this gets rid of extra shit at the ends of VCF names if it's included
		#	print(name, file=sys.stderr)
			name=name.replace(Trim,"") 
		#	print(name, Trim, "\n", file=sys.stderr)
		print(",", name, sep="", end="", file=OUT)
		names.append(name)
	print("\n", end="", file=OUT)
	return(cols)
	

	
		
	
def	format_Pheno(cols):
	#second line: individual numbers - just need something
	print("IndNum,", sep=",", end="", file=OUT)
	for i in range(0,cols):		
		print(",", i+1, sep="", end="", file=OUT)
	print("\n", end="", file=OUT)
	#print(names, file=sys.stderr)
	#now any more phenotype lines that there may be:
	if Pheno: #skip this bit if there is no phenotype file
		Pheno_dict={}
		#parse it into dictionaries for each phenotype: this dict has each ID as a key and then a list of phenos, the list of phenos will be in the same order/position as the phenoNames
		with open(Pheno, 'r') as PHENO:	
			for line in PHENO:
				if line.startswith("ID"):
					phenoNames=line.split()[1:]
				else:	
					fields=line.split()
					if fields[0] not in names:
						print("\nThere is a discrepancy between the names in the VCF and those in the phenotype file!", file=sys.stderr)
						print("VCF names look like this:", names[0], file=sys.stderr)
						print("Phenotype names look like this:", fields[0], file=sys.stderr)
						print("this can be addressed with the -t flag.", file=sys.stderr)
						print("It is also possible that ", fields[0], " is a parent or F1, in which case it may be ok to proceed.", sep="", end="\n", file=sys.stderr)
						try:
							continueOn=input_with_timeout("Proceed? y/n ", 15)
						except TimeoutExpired:
							print("  timeout expired, continuing on.", file=sys.stderr)
							continueOn="y"
						if continueOn != "y":	
							exit()
					else:
						Pheno_dict[fields[0]]=fields[1:]
		#now go through and print each phenotype based for each individual.
		for i in range(0,len(phenoNames)):
			print(phenoNames[i], end=",", file=OUT)
			for j in range(0,len(names)):
				print(",",Pheno_dict[names[j]][i], sep="",end="", file=OUT)
			print("\n", end="", file=OUT)
			
	


def format_Geno(str):
	fields=str.split("	")
#	print(fields, file=sys.stderr)
	ch=fields[0]
	if ch in ch_dict:
		ch_dict[ch]+=1
		if args.only_first_SNP:
			return(1) #this bit here forces the script to skip 2nd and 3rd... SNPs per each radtag - may be messing up later steps which try to find those steps. 
	else:
		ch_dict[ch]=1
	pos=fields[1]
	name=ch+"_"+pos
	print(name, "1", sep=",", end="", file=OUT)#the "1" was earlier 'ch' but the quide says make everything the same c'zome number if you don't have a map - though if you know the X linked ones, that's best...				
	j=0
	for fi in fields[9:]:
		j+=1
		if fi==".":
			f=fi
		else:		
			f=fi.split(":")[0]
		#print(fi)
		if f=="0/0":
			g="A"
		elif f=="0/1":	
			g="H"
		elif f=="1/1":
			g="B"
		elif f=="./.":
			g="NA"	
		elif f==".":
			g="NA"	
		else:
			g="UN"
		print(",", g, end="", sep="", file=OUT)
	print("\n", end="", sep="", file=OUT)				


ch_dict={}
with open(vcf, 'r') as VCF:
	for line in VCF:
		if not line.startswith("##"):
			if line.startswith("#"):
				cols=format_Header(line)		#first make the header row for the table: do this from the id's of the vcf file: ID,,ind1,ind2,ind3... followed by a row of sexes: sex,,1,1,1,2,2,1,1...
				format_Pheno(cols)			#second make the phenotype row for the table: do this from the Pheno dataset or lacking that, make a row of for each phenotype in the input file: pheno1,,x,y,z...	
			else:
				#print(line, file=OUT)
				format_Geno(line)		#third convert the vcf into genotypesA,H,B and NA: markerName,pos(1),A,B,H,NA,A...

 
 
 
```
