## Supplemental_File_5_Codebase for "A high-density genetic map and molecular sex-typing assay for gerbils": make_rqtl_input_file_for_Capture_het.py

```
#!/usr/bin/env python3

import re
import sys
import argparse
expl='''
make_rqtl_input_file_for_Capture_het.py
created by TDB 09-12-2014
updated for capture by TDB on 30-Aug-2016
updated again by TDB on 9-Feb-2018 to generate distance matrices rather than csvrs. It's not actually even for a capture anymore - it's to place the heterozygous markers on the map. 
	see 'make_rqlt_input_file.py' for the original. 


It generates two new matrices of the distances between the markers already on the map and the new markers to be placed on the map - one is the actual genotypes and one has the switched alleles. 
These matrices have individuals as columns (and have the individuals in the same order), though no colnames are present.
The one from the genetic map has RAD loci as rows while the one from the capture has SNPs as the rows.
The values in the body of the matrix are one for A, B, H, N where: A=AA hom, B=BB hom, H=AB het, N=anything else (NA)
The rownames are tab-separated from the rest of the table, but the values in the table are not separated by anything (will be a single string when imported into R)
'''
parser=argparse.ArgumentParser(description=expl)
parser.add_argument("--VCF", "-v", type=str, required=True, help="The VCF of F2s of markers that are heterozygous in the P generation - ES_Var.recode.vcf")
parser.add_argument("--VCFPandF1", "-w", type=str, required=True, help="The VCF parents and F1s of markers that are heterozygous in the P generation - PandF1.recode.vcf")
parser.add_argument("--Map", "-m", type=str, required=True, help="The Genetic map of RAD genotypes from gerbils - Meriones_map.csv perhaps")
parser.add_argument("--GenotypeQuality", "-GQ", type=int, required=False, help="Minimum genotype quality to output a genotype - otherwise genotype is NA", default=0)
parser.add_argument("--GenotypeDepth", "-GD", type=int, required=False, help="Minimum genotype depth required to output a genotype - otherwise genotype is NA", default=0)
parser.add_argument("--Completeness", "-c", type=int, required=False, help="Required completeness (number of genotypes NOT missing out of 137 gerbils) to test a loci against the rads", default=70)
parser.add_argument("--Out", "-o", type=str, required=True, help="Outfile name. This will be a matrix where rownames are RADs from the map (fixed in parents) and colnames are RADs that are heterozygous in the parents. Cell values are distances (0 is close, 137 is far). Omit for stdout")
#parser.add_argument("--CSVR", default=None, action='store_const', const=True, required=False, help="Forces script to make a csvr (lacking a header). Used to add genotypes to the map. Will be named after value declared by --out with .csvr as the stem.")

args=parser.parse_args()

vcf=args.VCF
vcfPandF1=args.VCFPandF1
map=args.Map
GQinput=args.GenotypeQuality
GDinput=args.GenotypeDepth
CompletenessInput=args.Completeness
out=args.Out

OUT=open(out, 'w')
OUT2=open(out.split(".")[0]+".SWITCHED_ALLELES.txt", 'w')
#if args.CSVR:
#	CSVR=sys.stdout#open(out.split(".")[0]+".csvr", 'w')


def format_name(name):
	fa=name.replace("_R1_clipped", "")
	fb=fa.rstrip("MF")
	#fc=fb.replace("BS.BB", "BSBB", 1)
	#f_out=fc.replace(".", "-")	
	return(fb)

def parse_geno(geno):
	fields=geno.split(":")
	#print(geno)#GT:DP:AD:GQ:GL
	geno=fields[0]
	if geno == "./.":
		AD="0,0"
		GD=0
		GQ=0
		PL="0,0,0"
	else:
		AD=fields[2]
		GD=int(fields[1])
		GQ=int(fields[3])
		PL=fields[4]
	if GQ < GQinput:									#here we can pick at cutoffs
		geno="./."
	elif GD < GDinput:
		geno="./."	
	#print(geno, file=sys.stderr)	
	g=format_geno(geno)
	return (g)

def format_geno(f):
	if f=="0/0" or f=="AA":
		g="A"		
	elif f=="0/1" or f=="AB" or f=="BA":
		g="H"
	elif f=="1/1" or f=="BB":
		g="B"
	elif f=="./." or f=="-":
		g="N"
	else:
		print("The genotypes cannot be formatted correctly:", f, file=sys.stderr)	
	return(g)

def switch_alleles(vec):#	
	#print(vec)
	switch_dict={"A":"B", "B":"A", "H":"H", "N":"N"}
	svec=[switch_dict[i] for i in vec]
	#print(svec)
	return(svec)

def calc_dist(vec1, vec2, toswitch="noSwitch", output="av"):
	#print(vec1, vec2, sep="\n", file=sys.stderr)
	if toswitch=="switch":
		vec1=switch_alleles(vec1)
	theSum=sum(1 for a,b in zip(vec1,vec2) if (a!=b and a!="N" and b!="N"))
	theLen=sum(1 for a,b in zip(vec1,vec2) if (a!="N" and b!="N"))
	out=theSum/theLen
	
	if output=="av":
		return(round(out,4)) #higher number = further apart
	elif output=="sum":
		return(theSum)
	
def get_c(geno1,geno2):
	C=geno1+geno2
	if re.search("N", C):#ungenotyped parent
		out=0
	elif C =="HH": #best - both F1s are hets
		out=2
	else: #one F1 is a het, the other is homozygous - this is possibly useable down the road. Ignoring it now.
		out=1
	return(out)

def get_useful_crosses(geneID,genos):
	#print(genos)
	P=genos[0]+genos[9]
	C1=get_c(genos[1],genos[4])
	C2=get_c(genos[2],genos[3])
	C3=get_c(genos[5],genos[8])
	C4=get_c(genos[6],genos[7])	

	if geneID not in useful_dict:
		useful_dict[geneID]={}
	useful_dict[geneID]["C1"]=C1 
	useful_dict[geneID]["C2"]=C2
	useful_dict[geneID]["C3"]=C3
	useful_dict[geneID]["C4"]=C4
	#now useful_dict can calculate back to whether a cross will have useful offspring -> 2=all offspring are useable!, 1=offspring mimic a backcross and probably only the hets should be used, 0=no offspring are usable from this cross. 
	
def drop_bad_inds(fields):
	#"ESES1-5M" and "ESES14-7F" were dropped form the map and shouldn't be included. 
	#print(len(fields), end="	")	
	del fields[35]  #index 36 is individual ESES14-7F - python base 0 means index 35
	del fields[4]	#index 5 is individual ESES1-5M   - python base 0 means index 4
	#print(len(fields), end="\n")
	return(fields)	
	
cross_origins={"ESES1-1":"C1", "ESES1-2":"C1", "ESES1-3":"C1", "ESES1-4":"C1", "ESES1-5":"C1", "ESES1-6":"C1", "ESES10-1":"C1", "ESES10-2":"C1", "ESES10-3":"C1", "ESES10-4":"C1", "ESES10-5":"C1", "ESES11-1":"C4", "ESES11-2":"C4", "ESES11-3":"C4", "ESES11-4":"C4", "ESES11-5":"C4", "ESES11-6":"C4", "ESES12-1":"C2", "ESES12-2":"C2", "ESES12-3":"C2", "ESES12-4":"C2", "ESES12-5":"C2", "ESES12-6":"C2", "ESES13-1":"C3", "ESES13-2":"C3", "ESES13-3":"C3", "ESES13-4":"C3", "ESES13-6":"C3", "ESES14-10":"C1", "ESES14-1":"C1", "ESES14-2":"C1", "ESES14-3":"C1", "ESES14-4":"C1", "ESES14-5":"C1", "ESES14-6":"C1", "ESES14-7":"C1", "ESES14-8":"C1", "ESES14-9":"C1", "ESES15-1":"C2", "ESES15-2":"C2", "ESES15-3":"C2", "ESES15-4":"C2", "ESES15-5":"C2", "ESES15-6":"C2", "ESES15-7":"C2", "ESES16-1":"C4", "ESES16-2":"C4", "ESES16-3":"C4", "ESES16-4":"C4", "ESES16-5":"C4", "ESES16-6":"C4", "ESES16-7":"C4", "ESES16-8":"C4", "ESES17-1":"C3", "ESES17-2":"C3", "ESES17-3":"C3", "ESES17-4":"C3", "ESES17-5":"C3", "ESES17-6":"C3", "ESES17-7":"C3", "ESES18-1":"C1", "ESES18-2":"C1", "ESES18-3":"C1", "ESES18-4":"C1", "ESES18-5":"C1", "ESES18-6":"C1", "ESES18-7":"C1", "ESES19-1":"C4", "ESES19-2":"C4", "ESES19-3":"C4", "ESES19-4":"C4", "ESES19-5":"C4", "ESES19-6":"C4", "ESES19-7":"C4", "ESES19-8":"C4", "ESES19-9":"C4", "ESES2-1":"C3", "ESES2-2":"C3", "ESES2-3":"C3", "ESES2-4":"C3", "ESES2-5":"C3", "ESES20-1":"C1", "ESES20-2":"C1", "ESES20-3":"C1", "ESES20-4":"C1", "ESES20-5":"C1", "ESES20-6":"C1", "ESES20-7":"C1", "ESES21-1":"C3", "ESES21-2":"C3", "ESES21-3":"C3", "ESES21-4":"C3", "ESES21-5":"C3", "ESES21-6":"C3", "ESES22-10":"C2", "ESES22-1":"C2", "ESES22-2":"C2", "ESES22-3":"C2", "ESES22-4":"C2", "ESES22-5":"C2", "ESES22-6":"C2", "ESES22-7":"C2", "ESES22-8":"C2", "ESES22-9":"C2", "ESES3-1":"C2", "ESES3-2":"C2", "ESES3-3":"C2", "ESES3-4":"C2", "ESES4-1":"C4", "ESES4-2":"C4", "ESES4-3":"C4", "ESES5-1":"C3", "ESES5-2":"C3", "ESES5-3":"C3", "ESES5-4":"C3", "ESES5-5":"C3", "ESES5b-1":"C4", "ESES5b-2":"C4", "ESES5b-3":"C4", "ESES6-1":"C1", "ESES6-2":"C1", "ESES6-3":"C1", "ESES6-4":"C1", "ESES6-5":"C1", "ESES6-6":"C1", "ESES6-7":"C1", "ESES7-1":"C3", "ESES7-2":"C3", "ESES7-3":"C3", "ESES7-4":"C3", "ESES8-1":"C2", "ESES8-2":"C2", "ESES9-1":"C4", "ESES9-2":"C4", "ESES9-3":"C4", "ESES9-4":"C4", "ESES9-5":"C4"}
	
		 
#here I go through the P and F1 data and find the sites that while heterozygous in the Parents, had the proper inheritance pattern to the F1s so that the F2s are usable:
#what I mean is:
#P:    AA x AB    #this parent cross can yield:
#F1:   AA or AB   #either of these F1s, but only AB F1s are informative. 

#and to make F2s, I crossed F1s which could have been any of these crosses:
#F1 cross 1:	AB x AB   BEST - this is great -go ahead with these, they will work exactly like the ones in the map.  
#F1 cross 2:	AA x AB or AB x AA  OK - these ones will probably work, but I need to think more about them - it's now a backcross scheme, not a F2 scheme which means they may not be immediately comparable to the rest of the map
										#probably what this means is that I can use some of the offspring, but not all - if the offspring are hets they are informative, if they are homozygous then throw them out.
#F1 cross 3:	AA x AA   BAD - unusable, these are not informative and need to be dropped. 


#here are the headers:
#col: 	9					10					11					12					13					14					15					16					17					18
#		EEEE0-1F_R1_clipped	EESS2-1F_R1_clipped	EESS2-3F_R1_clipped	EESS2-4M_R1_clipped	EESS2-6M_R1_clipped	EESS3-1F_R1_clipped	EESS3-2F_R1_clipped	EESS3-3M_R1_clipped	EESS3-4M_R1_clipped	SSSS0-1M_R1_clipped
#P are cols 9 and 18
#F1 crosses are:		VCFcol	genosCOl
	#EESS	2-1Fx2-6M	10x13	1x4			pg121	offspring:	ESES 1, 6, 10, 14, 18, 20
	#		2-3Fx2-4M	11x12	2x3			pg124				ESES 3, 8, 12, 15, 22
	#		3-1Fx3-4M	14x17	5x8			pg123				ESES 2, 5,  7, 13, 17, 21
	#		3-2Fx3-3M 	15x16	6x7			pg122				ESES 4, 5b, 9, 11, 16, 19


useful_dict={}


print("\nStarting to read VCF file:", vcfPandF1, file=sys.stderr)
print("This is to find useful genotypes based on the inheritance of heterozygosity from the parent generation.", file=sys.stderr)

with open(vcfPandF1, 'r') as VCF:
	for line in VCF:
		line=line.strip('\n')
		if line.startswith("#"):
			if not line.startswith("##"):
				#format headers here? Maybe extract individual ids?
				a=1
				#print(line, file=sys.stderr)
		else:	
			fields=line.split("	")
			gene_ID="_pos".join(fields[0:2])
			genotypes=fields[9:]
			genos=[parse_geno(genotype) for genotype in genotypes]
			get_useful_crosses(gene_ID, genos)
			#print(genos)

#print(useful_dict["10205_pos27"])
#print(useful_dict["115409_pos65"])
genes=[]	#a list of just the genes that have sufficient coverage
Geno_df=[] #used to store the list of genotypes
linenum=0
GeneID_dict={}
print("\nStarting to read VCF file:", vcf, file=sys.stderr)
print("	min GQ is", GQinput, file=sys.stderr)
print("	min GD is", GDinput, file=sys.stderr)
print("	genotypes needed in at least", CompletenessInput, "individuals to be evaluated", file=sys.stderr)

with open(vcf, 'r') as VCF:
	for line in VCF:
		line=line.strip("\n")
		if line.startswith("#"):
			names=line.split("	")[9:]
			GeneIDs=[format_name(i) for i in names]
			for i in GeneIDs:
				GeneID_dict[i]=""
		else:
			linenum=linenum+1
			print("		Working on gene:", linenum, end="\r", file=sys.stderr)
			fields=line.split("	")
			genotypes=fields[9:]
			#print(line, file=sys.stderr)
			#genos =[parse_geno(genotype) for genotype in genotypes]
			#print(genos)
			#new#
			genos=[]
			#print(line)
			#this next bit makes the genotype list - it goes through and if the F1s both had the right genotype, it allows it, otherwise it replaces the genotype with "N"			
			#print("genotypes:	  ", end="	")
			
			#this bit works right - TDB 13-08-2018
			for genotype,i in zip(genotypes,GeneIDs):
				#print(parse_geno(genotype), end=", ")
				#print(genotype, parse_geno(genotype), file=sys.stderr)
				if useful_dict["_pos".join(fields[0:2])][cross_origins[i]]==2:
					genos.append(parse_geno(genotype))	
				else:
					genos.append("N")	
			#print("\ngenoes		 	", ", ".join(genos), "\n")	
					
					
								
			#print(genos, "\n")
			completeness=sum(1 for i in genos if i!="N")
			#This will tell how many genes are actually useful. 
			#print(completeness, file=sys.stdout) #use this to generate the completeness.txt and completeness_hetF1xhomF1.txt files also make sure that the "=2" or "=1" above ~8 lines is set correctly

			if completeness >= CompletenessInput:
				genes.append("_pos".join(fields[0:2])) #list of gene names
				Geno_df.append(drop_bad_inds(genos))				   #list of genotypes for those gene names - list of lists. 
				
print("		Working on gene:", linenum, end="\n", file=sys.stderr)
print("	", len(genes), "genes entered with genotypes in >=", CompletenessInput,"individuals",  file=sys.stderr)

#print(Geno_df) # a list of lists. inner list is the genotypes of each gene
#print(genes) # a list of gene names, specifes the order of the genotypes. 
#print(genes[1:5])
#print(Geno_df[1:5])
#exit()

#for k in GeneID_dict:
#	print(k, file=sys.stderr)
print("Now reading in RADs from the genetic map:", map, "and comparing them with the VCF", file=sys.stderr)
print("Chr","cM", "	".join(genes) , sep="	", file=OUT)
print("Chr","cM", "	".join(genes) , sep="	", file=OUT2)

RADdict={}
linenum=0
outGenos=[]

with open(map, 'r') as MAP:
	for line in MAP:
		line=line.strip("\n")
		if line.startswith("ID"):
			fields=line.split(",")
			RADIDs=[format_name(i) for i in fields[3:]]
		elif line.startswith("NHTI") or line.startswith("NW_") or re.search("^\d", line): 
			linenum=linenum+1
			print("		Working on RAD:", linenum, end="\r", file=sys.stderr)
			fields=line.split(",")
			RAD=fields[0]
			chr=fields[1]
			cM=fields[2]
			loc=chr+"	"+cM
			
			mapgenos=[format_geno(i) for i in fields[3:]]
			#print(chr+"	"+cM,mapgenos, Geno_df[0], "\n", sep="\n")
			#print(len(mapgenos), len(Geno_df[0]))
			
			distVec=[calc_dist(hetgenos, mapgenos, toswitch="noSwitch", output="av") for hetgenos in Geno_df] #this compares the genotypes of map-markers with markers-to-add
			#for hetgenos in Geno_df:
			#	if chr == "13" or chr =="17" or chr =="19":
			#		print(chr, cM, calc_dist(hetgenos, mapgenos, toswitch="noSwitch", output="av"), sep="	", end="\n")
			
			
			print(loc, end="	", file=OUT)
			for i in distVec:
				print(i, end="	", file=OUT) 
			print("", file=OUT)	
			
			
			
			SwitchedAllele_distVec=[calc_dist(hetgenos, mapgenos, 'switch') for hetgenos in Geno_df]
			print(loc, end="	", file=OUT2)			
			for i in SwitchedAllele_distVec:
				print(i, end="	", file=OUT2)
			print("", file=OUT2)
			
			
print("		Working on RAD:", linenum, end="\r", file=sys.stderr)
print("", file=sys.stderr)

#print(RADIDs)
#print(len(outGenos[1]))
#print(len(GeneIDs))

#if args.CSVR:
#	print("TAG_ID","Chr", "cM", "	".join(GeneIDs), sep="	", file=CSVR)
#	for j in range(len(outGenos)):
#		print(allgenes[j],"NA", "NA", "	".join(outGenos[j]), sep="	", file=CSVR)


OUT.close()
OUT2.close()
#if args.CSVR:
#	CSVR.close()
```
