## Supplemental_File_5_Codebase for "A high-density genetic map and molecular sex-typing assay for gerbils": Meriones_Map_Het_markers_rqtl.r

```
#!/usr/local/bin/Rscript
library(qtl)
setwd("/Volumes/LaCie/Projects/Meriones_Map/stacks_out")

MerMap<-read.cross(format="csvr", file="Meriones_Map_AllMarkers.csv", na.strings=c("UN","NA","-"), dir="~/Documents/Meriones_Map/stacks_out", estimate.map=T, genotypes=c("AA", "AB", "BB"), alleles=c("A", "B"), convertXdata=F); summary(MerMap) 
names(MerMap$geno)<-c(1:21,"X")


MerMap<-est.rf(MerMap, tol=1e-4, maxit=1000)


#most of the problems here are because of missing data - it looks like it isn't all AA, but it's cause only a couple individuals were sequenced at those markers. 
gt<-geno.table(MerMap, scanone.output=T)
par(mfrow=c(2,1))
plot(gt, ylab=expression(paste(-log[10], " P-Value")), ylim=c(0,20))
abline(h=5,lty=2, col="gray")
plot(gt, lod=3:5, ylab="Genotype frequency")
abline(h=c(0.25, 0.5), lty=2, col="gray")
todrop<-rownames(gt[gt$neglog10P>5,]);todrop<-todrop[!(todrop %in%markernames(MerMap, chr="X"))];length(todrop);todrop # so maybe a lot to drop...?

cAlleles<-checkAlleles(MerMap, threshold=5);cAlleles; nrow(cAlleles)
#toswitch<-cAlleles$marker
#MerMap<-switchAlleles(MerMap, toswitch)

MerMap<-calc.errorlod(MerMap, error.prob=0.005)
print(toperr<-top.errorlod(MerMap, cutoff=6))

MerMap.clean<-MerMap
for(i in 1:nrow(toperr)){
	chr<-toperr$chr[i]
	id<-toperr$id[i]
	mar<-toperr$marler[i]
	MerMap.clean$geno[[chr]]$data[MerMap$pheno$id==id, mar] <- NA
}


write.cross(MerMap.clean, format="csvr", filestem="Meriones_Map_AllMarkers_final", digits=1)
summaryMap(MerMap)


if(FALSE){

#scaffs<-read.table("Xscaffs_for_rqtl.txt", header=F)$V1
error=0.0075
plotMap(MerMap.clean)

par(mfrow=c(2,2), las=1);
plot(ntyped(MerMap, "ind"), ylab="No. typed markers", main="No. genotypes by individual", pch=19, cex=.4, ylim=c(0,4500)) ;
plot(ntyped(MerMap, "mar"), ylab="No. typed individuals",  main="No. genotypes by marker", ylim=c(0,150), pch=19, cex=.2)
plot(sort(ntyped(MerMap, "ind"), decreasing=T), ylab="No. typed markers", main="No. genotypes by individual sorted", pch=19, cex=.4, ylim=c(0,4500)) ;
plot(sort(ntyped(MerMap, "mar"), decreasing=T), ylab="No. typed individuals",  main="No. genotypes by marker sorted", ylim=c(0,150), pch=19, cex=.2)
dev.new()
plotMap(MerMap)
dev.new()

newmap<-est.map(MerMap.clean, error=error, maxit=10000, sex.sp=F, verbose=T, tol=1e-4)
MerMap.clean<-replacemap(MerMap.clean, newmap)
plotMap(MerMap.clean)
summaryMap(MerMap.clean)


#just take a quick look through each chr and see what they look like now:
chr="X"
plotRF(MerMap, chr=chr)
plotMap(MerMap, chr=chr)
plotMissing(MerMap, chr=chr)
#MerMap<-Ripple(MerMap, chr=chr, window=7)


chr=21
plotRF(MerMap, chr=chr)
plotMap(MerMap, chr=chr)
plotMissing(MerMap, chr=chr)

chr=20
plotRF(MerMap, chr=chr)
plotMap(MerMap, chr=chr)
plotMissing(MerMap, chr=chr)
#MerMap<-Ripple(MerMap, chr=chr, window=7)

chr=19
plotRF(MerMap, chr=chr)
plotMap(MerMap, chr=chr)
plotMissing(MerMap, chr=chr)
#MerMap<-Ripple(MerMap, chr=chr, window=7)

chr=17
plotRF(MerMap, chr=chr)
plotMap(MerMap, chr=chr)
plotMissing(MerMap, chr=chr)
#MerMap<-Ripple(MerMap, chr=chr, window=7)

chr=16
plotRF(MerMap, chr=chr)
plotMap(MerMap, chr=chr)
plotMissing(MerMap, chr=chr)
#MerMap<-Ripple(MerMap, chr=chr, window=7)

chr=15
plotRF(MerMap, chr=chr)
plotMap(MerMap, chr=chr)
plotMissing(MerMap, chr=chr)
#MerMap<-Ripple(MerMap, chr=chr, window=7)

chr=14
plotRF(MerMap, chr=chr)
plotMap(MerMap, chr=chr)
plotMissing(MerMap, chr=chr)
#MerMap<-Ripple(MerMap, chr=chr, window=7)

chr=13
plotRF(MerMap, chr=chr)
plotMap(MerMap, chr=chr)
plotMissing(MerMap, chr=chr)
#MerMap<-Ripple(MerMap, chr=chr, window=7)

chr=12
plotRF(MerMap, chr=chr)
plotMap(MerMap, chr=chr)
plotMissing(MerMap, chr=chr)
#MerMap<-Ripple(MerMap, chr=chr, window=7)

chr=11
plotRF(MerMap, chr=chr)
plotMap(MerMap, chr=chr)
plotMissing(MerMap, chr=chr)
#MerMap<-Ripple(MerMap, chr=chr, window=7)

chr=10
plotRF(MerMap, chr=chr)
plotMap(MerMap, chr=chr)
plotMissing(MerMap, chr=chr)
#MerMap<-Ripple(MerMap, chr=chr, window=7)

chr=9
plotRF(MerMap, chr=chr)
plotMap(MerMap, chr=chr)
plotMissing(MerMap, chr=chr)
#MerMap<-Ripple(MerMap, chr=chr, window=7)

chr=8
plotRF(MerMap, chr=chr)
plotMap(MerMap, chr=chr)
plotMissing(MerMap, chr=chr)
#MerMap<-Ripple(MerMap, chr=chr, window=7)

chr=7
plotRF(MerMap, chr=chr)
plotMap(MerMap, chr=chr)
plotMissing(MerMap, chr=chr)
#MerMap<-Ripple(MerMap, chr=chr, window=7)

chr=6
plotRF(MerMap, chr=chr)
plotMap(MerMap, chr=chr)
plotMissing(MerMap, chr=chr)
#MerMap<-Ripple(MerMap, chr=chr, window=7)

chr=5
plotRF(MerMap, chr=chr)
plotMap(MerMap, chr=chr)
plotMissing(MerMap, chr=chr)
#MerMap<-Ripple(MerMap, chr=chr, window=5)

chr=4
plotRF(MerMap, chr=chr)
plotMap(MerMap, chr=chr)
plotMissing(MerMap, chr=chr)
#MerMap<-Ripple(MerMap, chr=chr, window=7)

chr=3
plotRF(MerMap, chr=chr)
plotMap(MerMap, chr=chr)
plotMissing(MerMap, chr=chr)
#MerMap<-Ripple(MerMap, chr=chr, window=7)

chr=2
plotRF(MerMap, chr=chr)
plotMap(MerMap, chr=chr)
plotMissing(MerMap, chr=chr)
#MerMap<-Ripple(MerMap, chr=chr, window=7)

chr=1
plotRF(MerMap, chr=chr)
plotMap(MerMap, chr=chr)
plotMissing(MerMap, chr=chr)
#MerMap<-Ripple(MerMap, chr=chr, window=7)

plotMap(MerMap)
}
```
