## Supplemental_File_5_Codebase for "A high-density genetic map and molecular sex-typing assay for gerbils": Meriones_Map_rqtl2.R

```
#this script was written by TDB in to creat the genetic map for meriones gerbils. 
#this is the final version


library(qtl)
setwd("/Volumes/LaCie/Projects/Meriones_Map/stacks_out")
MerMap<-read.cross(format="csvr", file="rqtl_input_2.csvr", na.strings=c("UN","NA"), dir="/Volumes/LaCie/Projects/Meriones_Map/stacks_out", estimate.map=FALSE); summary(MerMap) 

scaffs<-read.table("Xscaffs_for_rqtl.txt", header=F, stringsAsFactors=F)$V1 # problem - this set of markers has the genome scaffold names attached, but the scaffolds are to teh NHTI genome (GCA) rather than the NW genome (GCF). The GCF genome has the genes annotated and so it's better to use that one, but it wasn't around when I started the project which is why the NHTI names are used here, 
#so instead, for now, here are the important X scaffolds that will be needed with the appropriate scaffold nomenclature:
scaffs2<-c("NW_018668062.1_17124_rad_4334_244", "NW_018672145.1_4616_rad_74030_51", "NW_018661062.1_68605_rad_172011_190")

Ripple<-function(object, chr, window=4, method="countxo", error.prob=0.0001, breakAt=10){
	rip<-ripple(object, chr=chr, window=window, method=method, error.prob=error.prob)
	a=1
	while(as.data.frame(summary(rip))$obligXO[1]!=as.data.frame(summary(rip))$obligXO[2]){
		cat("switching order", a,"\n")
		object<-switch.order(object, chr=chr, rip[2,])
		rip<-ripple(object, chr=chr, window=window, method=method,error.prob=error.prob)
		a=a+1
		if(a>breakAt){cat("It doesn't seem to be converging...\n");break}
	}
	return(object)
}

mergeCHR<-function(object, chr1, chr2, switch=T){
	if(!(chr1 %in% names(object$geno)) | !(chr2 %in% names(object$geno))){
		cat("\n\n!!one of the chrs doesn't exist - try again!!\n\n")
		return(object)
	}
	chrSmall<-min(chr1,chr2)
	chrBig<-max(chr1,chr2)
	i=500
	markers<-markernames(object, chrBig)
	if(switch){object<-switchAlleles(object, markers)}
	for (marker in markers){
		i=i+1
		object<-movemarker(object, marker, chrSmall, i)		
	}	
	return(object)
}


nt.byind <-ntyped(MerMap, "ind")
nt.bymar <- ntyped(MerMap, "mar")

indstodrop<- c("ESES1-5M", "ESES14-7F") #names(nt.byind[nt.byind < 2000]); length(indstodrop)#this gets rid of markers genotyped in less than 60% of individuals - pretty strict, but a good idea. bad inds are:
indstokeep<-names(nt.byind)[!(names(nt.byind)%in%(indstodrop))];length(indstokeep)
MerMap<-subset(MerMap, ind=indstokeep);summary(MerMap)

todrop <- names(nt.bymar[nt.bymar < .84*135]);length(todrop);tokeep<-names(nt.bymar);tokeep<-tokeep[!(tokeep%in%todrop)];length(tokeep);todrop<-todrop[!(todrop%in%scaffs)];length(todrop) # 82% keeps in 8 X-linked scaffs, 85 keeps in 2, 84 keeps in 4
MerMap <- drop.markers(MerMap, todrop);marsleft=length(markernames(MerMap));marsleft;par(mfrow=c(1,2), las=1);plot(ntyped(MerMap, "ind"), ylab="No. typed markers", main="No. genotypes by individual", pch=19, cex=.4, ylim=c(0,marsleft)) ;plot(ntyped(MerMap, "mar"), ylab="No. typed individuals",  main="No. genotypes by marker", ylim=c(0,150), pch=19, cex=.2); summary(MerMap);summaryMap(MerMap)

par(mfrow=c(2,2), las=1);
plot(ntyped(MerMap, "ind"), ylab="No. typed markers", main="No. genotypes by individual", pch=19, cex=.4, ylim=c(0,marsleft)) ;
plot(ntyped(MerMap, "mar"), ylab="No. typed individuals",  main="No. genotypes by marker", ylim=c(0,150), pch=19, cex=.2)
plot(sort(ntyped(MerMap, "ind"), decreasing=T), ylab="No. typed markers", main="No. genotypes by individual sorted", pch=19, cex=.4, ylim=c(0,marsleft)) ;
plot(sort(ntyped(MerMap, "mar"), decreasing=T), ylab="No. typed individuals",  main="No. genotypes by marker sorted", ylim=c(0,150), pch=19, cex=.2)

gt<-geno.table(MerMap)
gt[gt$P.value <0.05/totmar(MerMap),]
todrop<-rownames(gt[gt$P.value <0.05/totmar(MerMap),]);todrop<-todrop[!(todrop%in%scaffs)];length(todrop) #
MerMap <- drop.markers(MerMap, todrop)

print(dup<-findDupMarkers(MerMap, exact.only=F));length(dup) #the number of duplicate markers - 60 groups comprising 129 markers - this isn't a lot... get rid of them.
length(unlist(dup))
todrop<-unlist(dup);length(todrop)
MerMap<-drop.markers(MerMap, todrop)
summary(MerMap)

MerMap <- est.rf(MerMap)
rf <- pull.rf(MerMap)
lod <- pull.rf(MerMap, what="lod")
dev.new();plot(as.numeric(rf), as.numeric(lod), xlab="Recombination fraction", ylab="LOD score", pch=20, cex=.4);dev.new()


backup_map1<-MerMap
MerMap<-backup_map1
RF=0.3;   
LOD=4;    
lg <- formLinkageGroups(MerMap, max.rf=RF, min.lod=LOD); table(lg[,2])

MerMap<-formLinkageGroups(MerMap, max.rf=RF, min.lod=LOD, reorgMarkers=T) # this makes 51 linkage groups, but it's probably because the alleles are switched in some and so everything is broken into two.
MerMap <- est.rf(MerMap)
plotRF(MerMap, chr=c(1:51), alt=T)

MerMap<-mergeCHR(MerMap, 1,29);#plotRF(MerMap, alternate=T)
MerMap<-mergeCHR(MerMap, 2,41);#plotRF(MerMap, alternate=T)
MerMap<-mergeCHR(MerMap, 3,11);#plotRF(MerMap, alternate=T)
MerMap<-mergeCHR(MerMap, 4,47);#plotRF(MerMap, alternate=T)
MerMap<-mergeCHR(MerMap, 5,20);#plotRF(MerMap, alternate=T)
MerMap<-mergeCHR(MerMap, 6,44);#plotRF(MerMap, alternate=T)
MerMap<-mergeCHR(MerMap, 7,36);#plotRF(MerMap, alternate=T)
MerMap<-mergeCHR(MerMap, 8,43);#plotRF(MerMap, alternate=T)
MerMap<-mergeCHR(MerMap, 8,50);#plotRF(MerMap, alternate=T)
MerMap<-mergeCHR(MerMap, 9,42);#plotRF(MerMap, alternate=T)
MerMap<-mergeCHR(MerMap, 10,35);#plotRF(MerMap, chr=c(10:51),alternate=T)
MerMap<-mergeCHR(MerMap, 13,21);#plotRF(MerMap, chr=c(13:51),alternate=T)
MerMap<-mergeCHR(MerMap, 14,46);#plotRF(MerMap, chr=c(14:51),alternate=T)
MerMap<-mergeCHR(MerMap, 15,18);#plotRF(MerMap, chr=c(15:51),alternate=T)
MerMap<-mergeCHR(MerMap, 16,39);#plotRF(MerMap, chr=c(16:51),alternate=T)
MerMap<-mergeCHR(MerMap, 17,48);#plotRF(MerMap, chr=c(17:51),alternate=T)
MerMap<-mergeCHR(MerMap, 19,27);#plotRF(MerMap,chr=c(19:51), alternate=T)
MerMap<-mergeCHR(MerMap, 23,25);#plotRF(MerMap, chr=c(23:51),alternate=T)
MerMap<-mergeCHR(MerMap, 24,32);#plotRF(MerMap, chr=c(24:51),alternate=T)
MerMap<-mergeCHR(MerMap, 26,38);#plotRF(MerMap, alternate=T)
MerMap<-mergeCHR(MerMap, 28,40);#plotRF(MerMap, alternate=T)
MerMap<-mergeCHR(MerMap, 30,37);#plotRF(MerMap, alternate=T)
MerMap<-mergeCHR(MerMap, 45,51);#plotRF(MerMap, alternate=T)

backup_map2<-MerMap
MerMap<-backup_map2
plotMap(MerMap)
chr=18


MerMap <- est.rf(MerMap)

for (chr in names(MerMap$geno)){
	cat("working on chr", chr, "\n")
	dev.new()
	par(mfrow=c(1,2))
	plotRF(MerMap, chr=chr, alternate.chrid=T)
	MerMap<-orderMarkers(MerMap, chr=chr, use.ripple=T, window=4, sex.sp=F, verbose=T)
	plotRF(MerMap, chr=chr, alternate.chrid=T)
}
dev.new()
plotRF(MerMap, alternate.chrid=T)
plotMap(MerMap)

MerMap <- est.rf(MerMap)
#to re-run of just some specific ones until things look good
if(FALSE){
	for (chr in c(15,10,5,2)){
		cat("working on chr", chr, "\n")
		dev.new()
		par(mfrow=c(1,2))
		plotRF(MerMap, chr=chr, alternate.chrid=T)
		MerMap<-orderMarkers(MerMap, chr=chr, use.ripple=T, window=4, sex.sp=F, verbose=T)
		plotRF(MerMap, chr=chr, alternate.chrid=T)
	}
}

backup_map3<-MerMap
MerMap<-backup_map3

plotMap(MerMap)


#finally, re-order based on new size:
MerMap$geno<-MerMap$geno[order(summaryMap(MerMap)[1:28,2], decreasing=T)]
names(MerMap$geno)<-c(1:28)
MerMap <- est.rf(MerMap)

backup_map4<-MerMap
MerMap<-backup_map4


#find the X:
gt<-geno.table(MerMap, scanone.output=T)
par(mfrow=c(2,1))
plot(gt, ylab=expression(paste(-log[10], " P-Value")), ylim=c(0,10))
abline(h=5,lty=2, col="gray")
plot(gt, lod=3:5, ylab="Genotype frequency")
abline(h=c(0.25, 0.5), lty=2, col="gray")
#put it at the end:
MerMap$geno<-MerMap$geno[c(1:17,19:28,18)]
plotMap(MerMap)
names(MerMap$geno)<-c(1:27,"X")
MerMap <- est.rf(MerMap)

summary(MerMap)


backup_map5<-MerMap
MerMap<-backup_map5

#Any more merges? 
par(mfrow=c(1,1))
plotRF(MerMap, col="redblue")

#this one seems most likely
MerMap<-mergeCHR(MerMap, 7,22, switch=T);plotRF(MerMap, chr=7, col="redblue")
MerMap<-orderMarkers(MerMap, chr=7, use.ripple=T, window=5, sex.sp=F, verbose=T);MerMap<-est.rf(MerMap);plotRF(MerMap, chr=7)
MerMap<-Ripple(MerMap, chr=7, window=7)

MerMap<-mergeCHR(MerMap, 11,24, switch=T);plotRF(MerMap, chr=11, col="redblue")
MerMap<-orderMarkers(MerMap, chr=11, use.ripple=T, window=5, sex.sp=F, verbose=T);MerMap<-est.rf(MerMap);plotRF(MerMap, chr=11)
compareorder(MerMap, chr=11, order=c(22:28,1:21))
MerMap<-switch.order(MerMap, chr=11, c(22:28,1:21))
plotRF(MerMap, chr=11, col="redblue")
MerMap<-Ripple(MerMap, chr=11, window=7)
plotRF(MerMap, chr=11, col="redblue")


plotRF(MerMap, col="redblue")
plotMap(MerMap)
summaryMap(MerMap)


#get rid of small chrs and ones with a single marker:
todrop<-markernames(MerMap, chr=c(1,27,26,25))
MerMap <- drop.markers(MerMap, todrop)

#re-order and name properly:
MerMap$geno<-MerMap$geno[order(summaryMap(MerMap)[1:22,2], decreasing=T)]
MerMap$geno<-MerMap$geno[c(1:16,18:22,17)]
names(MerMap$geno)<-c(1:21,"X")
plotMap(MerMap)
MerMap <- est.rf(MerMap)
plotRF(MerMap, col="redblue", alt=T)


MerMap<-est.rf(MerMap)


for (chr in names(MerMap$geno)){
	cat("working on chr:", chr, "\n")
	MerMap<-Ripple(MerMap, chr=chr, window=7)
}

plotRF(MerMap, chr=1, col="redblue")

compareorder(MerMap, chr=1, order=c(39:57, 1:38))
MerMap<-switch.order(MerMap, chr=1, order=c(39:57,1:38))
MerMap<-Ripple(MerMap, chr=1, window=7)


dropone<-droponemarker(MerMap, sex.sp=F)
par(mfrow=c(2,1))
plot(dropone, lod=1, ylim=c(-100,0))
plot(dropone, lod=2, ylab="Change in chromosome length")
par(mfrow=c(1,1))
dropone[dropone$Ldiff>10,] #markers that change the length a lot - perhaps drop them out.
print(todrop<-rownames(dropone[dropone$Ldiff>10,]))
#they almost all look like terminal markers so don't drop them! BUT There is one that is a middle marker on chr 1:
todrop<-todrop[todrop %in%markernames(MerMap, chr=1)];length(todrop)
MerMap<-drop.markers(MerMap,todrop)
summaryMap(MerMap)
#now re-estimate map:
MerMap <- est.rf(MerMap)
newmap<-est.map(MerMap, sex.sp=F)
MerMap<-replace.map(MerMap, newmap)
summaryMap(MerMap)
plotMap(MerMap)
plotRF(MerMap)

if (FALSE){#this takes a bit of time, so to save time I don't run it always
	loglik<-err<-c(0.0010, 0.0025, 0.0050, 0.0075, 0.0100, 0.0125, 0.0150, 0.0175, 0.0200) 
	for (i in seq(along=err)){
		cat(i, "of", length(err), "	")
		tempmap<-est.map(MerMap, error.prob=err[i], tol=1e-4)
		loglik[i]<-sum(sapply(tempmap, attr, "loglik"))
		cat("loglik is:", loglik[i], "for err:", err[i], "\n")
	}
	lod<-(loglik - max(loglik))/log(10)
	plot(err, lod, xlab="Genotyping error rate", ylab=expression(paste(log[10], " likelihood")), pch=20)
	error=err[lod==max(lod)]
}

error=0.0075 
save.image(file="Meriones_Map2_1.RData")

################################################################################################
#phase 2
################################################################################################
library(qtl)
setwd("~/Documents/Meriones_Map/stacks_out")
load("Meriones_Map2_1.RData")

#write.cross(format="mapqtl", cross=MerMap, chr=1, filestem="MerMap_chr1")

dropone<-droponemarker(MerMap, sex.sp=F)
par(mfrow=c(2,1))
plot(dropone, lod=1, ylim=c(-100,0))
plot(dropone, lod=2, ylab="Change in chromosome length")
par(mfrow=c(1,1))
dropone[dropone$Ldiff>10,] #markers that change the length a lot - perhaps drop them out.
print(todrop<-rownames(dropone[dropone$Ldiff>10,]))
#they all look like terminal markers so don't drop them! 
#MerMap<-drop.markers(MerMap,todrop)
summaryMap(MerMap)


plot(countXO(MerMap), ylab="Number of Crossovers", pch=21, bg="red", xlab="Individual")
MerMap<-calc.errorlod(MerMap, error.prob=error)
allerr<-top.errorlod(MerMap, cutoff=0)
hist(allerr$errorlod, col="red", breaks=400)
head(toperr<-top.errorlod(MerMap, cutoff=1));nrow(toperr) #this many genotypes are possibly suspect - very high, but reasonable - many genotypes had really low qualities/coverage

#look at chrs and XOs
ind=c(1:80)
plotGeno(MerMap, chr=1,  ind=ind,cutoff=6, include.xo=T, min.sep=2)
summary(MerMap)
table(toperr$chr)


backup7<-MerMap
#get rid of those genotypes (also could correct them, but lets start this way)
for(i in seq(from=1, to=nrow(toperr))){
	chr<-toperr$chr[i]
	ind<-toperr$id[i]
	mar<-toperr$marker[i]
#	cat(chr, ind, mar, "\n")
	MerMap$geno[[chr]]$data[MerMap$pheno$ID==ind, mar]<-NA	
}
summaryMap(MerMap)


#look at some summary things:
plot(countXO(MerMap), ylab="Number of Crossovers", pch=21, bg="red", xlab="Individual")
plotGeno(MerMap, chr="X", ind=c(25:65),cutoff=10, include.xo=T, min.sep=2)
barplot(colSums(countXO(MerMap, bychr=T)), col="blue", main="number of XO's per chromosome")
i=1
hist(countXO(MerMap, bychr=T)[,i], breaks=0:10, col="blue", ylim=c(0,140), xlim=c(-1,12));i;i=i+1
#so of those histograms, they should all be centered at 1 with a variance of 1 (poisson distributed) - looks good


#now re-visit segregation distortion:
gt<-geno.table(MerMap, scanone.output=T)
par(mfrow=c(2,1))
plot(gt, ylab=expression(paste(-log[10], " P-Value")), ylim=c(0,10))
abline(h=5,lty=2, col="gray")
plot(gt, lod=3:5, ylab="Genotype frequency")
abline(h=c(0.25, 0.5), lty=2, col="gray")
todrop<-rownames(gt[gt$neglog10P>5,]);todrop<-todrop[!(todrop %in%scaffs)];length(todrop);todrop
#MerMap<-drop.markers(MerMap, todrop)
#no appreciable SD


#now re-order LGs again:
#it would be nice to re-order LGs by length
summaryMap(MerMap)
MerMap$geno<-MerMap$geno[order(summaryMap(MerMap)[1:22,2], decreasing=T)]
MerMap<-est.rf(MerMap)
names(MerMap$geno)<-c(1:16, "X", 18:22)
#put the x at the end:
MerMap$geno<-MerMap$geno[c(1:16,18:22,17)]
plotMap(MerMap, alt=T)


#So this all loosk really good:
save.image(file="Meriones_Map2_2.RData")
write.cross(MerMap, format="csvr", filestem="Meriones_Map")

load("Meriones_Map2_2.RData")


setwd("~/Dropbox/Post_Doc/Manuscripts/Meriones_Map/figures/ECRs_genome_figure/data")
#write.cross(MerMap, format="csvr", filestem="Meriones_Map", digits=2)

summaryMap(MerMap)
plotMap(MerMap)
```
