## Supplemental_File_5_Codebase for "A high-density genetic map and molecular sex-typing assay for gerbils": Meriones_Stacks_5.sh

```
#!/bin/bash

##3 differs from #2 in that it will be run locally on mwk16. No more deepthought/UDrive shit.
#updated 18-04-2018

#this is the script that runs the GBS pipeline for the Meriones. 
#written by TDB on 6 November 2017. 


	#original raw gzipped data should be in:	 /UDrive/CNS/ResearchData/BIOL-GROUP-Mulley/tom_brekke/raw/Meriones_GBS_LGC_1a/RE_processed/
	#original raw gzipped data should be in:	 /UDrive/CNS/ResearchData/BIOL-GROUP-Mulley/tom_brekke/raw/Meriones_GBS_LGC_1b/RE_processed/
	#original raw gzipped data should be in:	 /UDrive/CNS/ResearchData/BIOL-GROUP-Mulley/tom_brekke/raw/Meriones_GBS_LGC_2/AdapterClipped/  #didn't come RE_processed, not sure why... But it also looks like RE processing doesn't remove much, so it shouldn't be much of a problem. - any wonky reads won't form stacks and will be dropped later anyway.

		#some of the raw data needs to be manipulated - 
	#1b needs to be renamed - they included sample names in the titles. 
		#here is how I renamed them:
		#for files in folders:
			#for i in S*E*/*; do echo $i; j=${i/\/[[:digit:]]*-E//\E}; mv $i $j; done
		#for folders:
			#for i in S*E*; do echo $i; j=${i/_[[:digit:]]*-E/_E};  mv $i $j; done	
	#2 needs to be as well:
		#this renames the files
			#for i in Sample_[[:digit:]]_*/*; do echo $i; j=${i/\/[[:digit:]]_/\/}; echo $j; mv $i $j; done
			#for i in Sample_[[:digit:]][[:digit:]]_*/*; do echo $i; j=${i/\/[[:digit:]][[:digit:]]_/\/}; echo $j; mv $i $j;done
			#for i in Sample_[[:digit:]][[:digit:]]-*/*; do echo $i; j=${i/\/[[:digit:]][[:digit:]]-/\/}; echo $j; mv $i $j;done
			#for i in Sample_[[:digit:]][[:digit:]][[:digit:]]-*/*; do echo $i; j=${i/\/[[:digit:]][[:digit:]][[:digit:]]-/\/}; echo $j; mv $i $j;done
			
		#this renames the dirs
			#for i in Sample_[[:digit:]][[:digit:]][[:digit:]]-*; do echo $i; j=${i/Sample_[[:digit:]][[:digit:]][[:digit:]]-/Sample_};echo $j ;mv $i $j;done
			#for i in Sample_[[:digit:]][[:digit:]][[:digit:]]_*; do echo $i; j=${i/Sample_[[:digit:]][[:digit:]][[:digit:]]/Sample};echo $j ;mv $i $j;done
			#for i in Sample_[[:digit:]][[:digit:]]-*; do echo $i; j=${i/Sample_[[:digit:]][[:digit:]]-/Sample_};echo $j ;mv $i $j;done
			#for i in Sample_[[:digit:]][[:digit:]]_*; do echo $i; j=${i/Sample_[[:digit:]][[:digit:]]/Sample};echo $j ;mv $i $j;done
			#for i in Sample_[[:digit:]]_*; do echo $i; j=${i/Sample_[[:digit:]]/Sample};echo $j ;mv $i $j;done
		#then some files I missed - the '-' ones...:
			#for i in Sample_*/[[:digit:]][[:digit:]][[:digit:]]*; do echo $i; j=${i/\/[[:digit:]][[:digit:]][[:digit:]]_/\/}; echo $j; mv $i $j; done
		


################################################
###  SET THESE PRIOR TO RUNNING THE SCRIPT:  ###
################################################

#This is the location of the working directory:
homedir="/Users/bss81d/Documents/Meriones_Map"

#These are the location of the raw data - external HD called MULLEY_LAB_Data:
	raw_data_dir_1a="/Volumes/MULLEY_LAB_Data/Meriones_GBS_LGC_1a_30-Oct-2017/AdapterClipped"
	raw_data_dir_1b="/Volumes/MULLEY_LAB_Data/Meriones_GBS_LGC_1b_08-Jan-2018/AdapterClipped"
	raw_data_dir_2="/Volumes/MULLEY_LAB_Data/Meriones_GBS_LGC_2_03-Apr-2018/AdapterClipped"

#for the genome - used to link radtags to a genome contig.
GENOME_LOCATION_GCF="/Users/bss81d/Dropbox/Post_Doc/Genomes/Meriones_unguiculatus/Genome_Lucigen/GCF_002204375.1_MunDraft-v1.0/GCF_002204375.1_MunDraft-v1.0_genomic.fna"
#run_cmd.sh "bwa index $GENOME_LOCATION_GCF" $GENOME_LOCATION_GCF.amb

################################################
###                                         ####
################################################


#this checks that the file structure is in place
if [ ! -f $homedir ]
then
	mkdir $homedir
fi	
cd $homedir
if [ ! -d stacks_out ]
then 	
	mkdir stacks_out
	mkdir alignments
	mkdir het_markers
	mkdir snps
	mkdir map-genome
fi
if [ ! -d ./raw/ ]
then
	mkdir ./raw
	echo "Raw fastqs after being merged from AdapterClipped directory in the raw data file" > ./raw/README.txt
fi			

	
#this first bit takes the multiple different sequencing efforts and merges them all together. 
if [ ! -f ./stacks_out/process_radtags_finished.txt ]
then
	cd $raw_data_dir_1a 
	ID=1
	for i in Sample* #folder that each sequencing bit is in
	do 
		cd $i
		for j in *R1_clipped.fastq.bz2 #name of the sequencing reads
		do	
			echo ""
			echo $j
		#do some unpacking
			run_cmd.sh "bunzip2 -kc $j > $homedir/raw/${j/.bz2/}; touch $homedir/raw/${j/.bz2/.0.done}" $homedir/raw/${j/.bz2/.0.done} & 
			k=${j/R1/R2} #here is the 2nd read file - it should be decompressed as well. 
			run_cmd.sh "bunzip2 -kc $k > $homedir/raw/${k/.bz2/}; touch $homedir/raw/${k/.bz2/.0.done}" $homedir/raw/${k/.bz2/.0.done}
			wait
		#check if the sample has been sequenced a second time, if it has, merge fastq files	
			if [ -f $raw_data_dir_1b/$i/$j ] 
			then  
			#do some unpacking and merging of resequenced reads with original library prep. Don't worry about the fact they are on different lanes and may have different error profiles - too complicated. 
				run_cmd.sh "bunzip2 -kc $raw_data_dir_1b/$i/$j >> $homedir/raw/${j/.bz2/}; touch $homedir/raw/${j/.bz2/.1.done}" $homedir/raw/${j/.bz2/.1.done} &
				run_cmd.sh "bunzip2 -kc $raw_data_dir_1b/$i/$k >> $homedir/raw/${k/.bz2/}; touch $homedir/raw/${k/.bz2/.1.done}" $homedir/raw/${k/.bz2/.1.done} 
				wait
			fi		
		#this bit should get the round 2 sequences together - unfortunately, some are with - and some with _ (for the samples that were sequenced 2x...)
		#will need to deal with that first...
		#set the folder - $i is sometimes found in batch 2, b2i is always found in batch 2  
		#	echo $i #sometimes will be found
			b2i2=${i/-/.} #always will be found
		#set the file $j - This is complicated since in the second batch of sequencing some are named with a "-" and some are named with an "_" - at least for the ones sequenced twice in batch2
			b2j=$j #new with just the Adapter_clipped
			b2j2=${b2j/-/.}				#for the ones with "." ->all of them
			b2k=${b2j/R1/R2}
			b2k2=${b2j2/R1/R2}

			#only some of them were sequenced 2x - these have $i and b2j
 			if [ -f $raw_data_dir_2/$i/$b2j ]
			then
				run_cmd.sh "bunzip2 -kc $raw_data_dir_2/$i/$b2j >> $homedir/raw/${j/.bz2/}; touch $homedir/raw/${j/.bz2/.2.done}" $homedir/raw/${j/.bz2/.2.done} &
				run_cmd.sh "bunzip2 -kc $raw_data_dir_2/$i/$b2k >> $homedir/raw/${k/.bz2/}; touch $homedir/raw/${k/.bz2/.2.done}" $homedir/raw/${k/.bz2/.2.done} 
				wait
 			fi
			#all of them were sequenced 1+ and these have $b2i2 and $b2j2
			run_cmd.sh "bunzip2 -kc $raw_data_dir_2/$b2i2/$b2j2 >> $homedir/raw/${j/.bz2/}; touch $homedir/raw/${j/.bz2/.3.done}" $homedir/raw/${j/.bz2/.3.done} &
			run_cmd.sh "bunzip2 -kc $raw_data_dir_2/$b2i2/$b2k2 >> $homedir/raw/${k/.bz2/}; touch $homedir/raw/${k/.bz2/.3.done}" $homedir/raw/${k/.bz2/.3.done} 
			wait
			cd $homedir/raw
			run_cmd.sh "process_radtags -i fastq -1 ${j/.bz2/} -2 ${k/.bz2/} -o ../stacks_out/ -q -y gzfastq -t 140 --disable_rad_check --len_limit 140 -c" ../stacks_out/${j/fastq.bz2/1.fq.gz}
			run_cmd.sh "mv ../stacks_out/process_radtags.log ../stacks_out/${j/.fastq.bz2/.log}" ../stacks_out/${j/.fastq.bz2/.log} 


		#now that all the reads are concatenated, it's time to start the actual stacks pipeline.


		#only run ustacks on the first reads though. and it was renamed earlier, so make sure to test for the absense of that too:
			if [ ! -f ../stacks_out/${j/fastq.bz2/1.snps.tsv.gz} ]
			then		
				run_cmd.sh "ustacks -f ../stacks_out/${j/fastq.bz2/1.fq.gz} -i $ID -o ../stacks_out/ -m 3 -M 2 -N 4 -H -p 3 -d --max_locus_stacks 3 --model_type snp --alpha 0.01" ../stacks_out/${j/fastq.bz2/snps.tsv.gz}		
			fi
			wait
			let "ID++"		
		done
		#echo "is this file here: $homedir/stacks_out/${j/fastq.bz2/snps.tsv.gz}" 
		cd $homedir/raw/
		cmd="rm $homedir/raw/${j/.bz2/}"
		echo "CMD: $cmd"
		cmd2="rm $homedir/raw/${k/.bz2/}"
		if [ -f $homedir/stacks_out/${j/fastq.bz2/1.snps.tsv.gz} ]		
		then
			echo "	evaluating"
			eval $cmd
			echo "CMD: $cmd2"
			echo "	evaluating"
			eval $cmd2
		fi
		cd $raw_data_dir_1a 
	done		
	touch $homedir/stacks_out/process_radtags_finished.txt
fi

cd $homedir


echo "finished process_radtags and ustacks for all individuals, "
cd ./stacks_out/

#need to do some renaming to get tsv2bam to work right:  R2 files have both an R2 and the .2. and so won't be recognised - name must match exactly except for the .1. or .2. - this changes all to "R1", but leaves the .1. and .2.
if [ ! -f renaming_finished.txt ]
then
	for i in *R2*  #this renames the 2nd read files so that they have R1 not R2, and read number is demarcated by the .1. or .2. at the end instead
	do
		j=${i/R2/R1}
		run_cmd.sh "mv $i $j" $j
	done	
	for i in *.1.*.tsv.gz  #this gets rid of the .1. and .2. bits in the files where they are not needed - the 'matches' and 'alleles' etc.
	do
		j=${i/.1./.}
		run_cmd.sh "mv $i $j" $j
	done	
	touch renaming_finished.txt 
fi
#now back to the stacks pipeline:
run_cmd.sh "cstacks -s EEEE0-1F_R1_clipped -s SSSS0-1M_R1_clipped -o ./ -p 4 -n 2 " catalog.tags.tsv.gz
run_cmd.sh "sstacks -P ./ -M ./population_map.txt -p 3; touch sstacks_finished.txt" sstacks_finished.txt
run_cmd.sh "tsv2bam -P ./ -R ./ -t 3 -M population_map.txt; touch tsv2bam_finished.txt" tsv2bam_finished.txt
run_cmd.sh "gstacks -P ./ -t 3 -M population_map.txt" gstacks.log 
run_cmd.sh "populations -V catalog.calls -O ./ -t 2 -p 1 --min_maf 0.01 --write_random_snp --vcf" catalog.calls.p.snps.vcf # the write_random_snp bit doesn't seem to work... not sure why. Dealt with the issue during "make_rqtl_input_file.py" which only uses one SNP per tag.


#pull out just the SNPs that are alternatively fixed in the parents as well as ones that are het in one parent but not the other:
run_cmd.sh "filter_SNPs.py -v catalog.calls.p.snps.vcf -o1 ES_Fixed.vcf -o2 ES_Var.vcf" ES_Fixed.vcf
#pull the parents and F1s out of the VCF so that it is easier to deal with in R/qtl (this may not be a good idea in the end)
#should consider some genotype filtering with the vcftools call here: --minGQ 20 --minDP 10
run_cmd.sh "vcftools --vcf ES_Fixed.vcf  --out ES_Fixed --recode --remove-indv EEEE0-1F_R1_clipped --remove-indv EESS2-1F_R1_clipped --remove-indv EESS2-3F_R1_clipped --remove-indv EESS2-4M_R1_clipped --remove-indv EESS2-6M_R1_clipped --remove-indv EESS3-1F_R1_clipped --remove-indv EESS3-2F_R1_clipped --remove-indv EESS3-3M_R1_clipped --remove-indv EESS3-4M_R1_clipped --remove-indv SSSS0-1M_R1_clipped" ES_Fixed.recode.vcf
#convert the VCF into a csvr for r/qtl.
run_cmd.sh "make_rqtl_input_file.py -v ES_Fixed.recode.vcf -t \"_R1_clipped\" -p ../Phenotype_data.txt -o rqtl_input.csvr --only_first_SNP" rqtl_input.csvr # this takes only the first SNP per radtag to build the Map 

#this bit associates the scaffolds with the markers that are fixed in each parent:
	#I should have used the other genomes GCF instead of GCA...
run_cmd.sh "awk -F '_'  'NR>7 {print \">\"\$1\" \"}' rqtl_input.csvr > genotyped_radtags_togrep.txt" genotyped_radtags_togrep.txt
run_cmd.sh "gunzip -c catalog.fa.gz > catalog.fa" catalog.fa
run_cmd.sh "extract_fasta_region.py -f catalog.fa -chrList genotyped_radtags_togrep.txt -fmt 10000 -o genotyped_radtags.fasta" genotyped_radtags.fasta
run_cmd.sh "bwa mem $GENOME_LOCATION_GCF genotyped_radtags.fasta > genotyped_radtags.sam" genotyped_radtags.sam
run_cmd.sh "rename_tags.py -s genotyped_radtags.sam -r rqtl_input.csvr > rqtl_input_2.csvr" rqtl_input_2.csvr 


#this does the same with markers that were het in one parent: consider the genotype filtering - --minGQ 20 --minGD 10
run_cmd.sh "vcftools --vcf ES_Var.vcf --out ES_Var --recode --remove-indv EEEE0-1F_R1_clipped --remove-indv EESS2-1F_R1_clipped --remove-indv EESS2-3F_R1_clipped --remove-indv EESS2-4M_R1_clipped --remove-indv EESS2-6M_R1_clipped --remove-indv EESS3-1F_R1_clipped --remove-indv EESS3-2F_R1_clipped --remove-indv EESS3-3M_R1_clipped --remove-indv EESS3-4M_R1_clipped --remove-indv SSSS0-1M_R1_clipped" ES_Var.recode.vcf
run_cmd.sh "vcftools --vcf ES_Var.vcf --out PandF1 --recode --indv EEEE0-1F_R1_clipped --indv EESS2-1F_R1_clipped --indv EESS2-3F_R1_clipped --indv EESS2-4M_R1_clipped --indv EESS2-6M_R1_clipped --indv EESS3-1F_R1_clipped --indv EESS3-2F_R1_clipped --indv EESS3-3M_R1_clipped --indv EESS3-4M_R1_clipped --indv SSSS0-1M_R1_clipped" PandF1.recode.vcf

run_cmd.sh "make_rqtl_input_file.py -v ES_Var.recode.vcf -t \"_R1_clipped\" -p ../Phenotype_data.txt -o rqtl_input_Var.csvr " rqtl_input_Var.csvr
run_cmd.sh "awk -F '_'  'NR>7 {print \">\"\$1\" \"}' rqtl_input_Var.csvr > genotyped_radtags_togrep_Var.txt" genotyped_radtags_togrep_Var.txt
run_cmd.sh "gunzip -c catalog.fa.gz > catalog.fa" catalog.fa
run_cmd.sh "extract_fasta_region.py -f catalog.fa -chrList genotyped_radtags_togrep_Var.txt -fmt 10000 -o genotyped_radtags_Var.fasta" genotyped_radtags_Var.fasta
run_cmd.sh "bwa mem $GENOME_LOCATION_GCF genotyped_radtags_Var.fasta > genotyped_radtags_Var.sam" genotyped_radtags_Var.sam
run_cmd.sh "rename_tags.py -s genotyped_radtags_Var.sam -r rqtl_input_Var.csvr> rqtl_input_Var_2.csvr" rqtl_input_Var_2.csvr

#now need to format the reference fasta correctly and index it:
run_cmd.sh "bwa index catalog.fa" catalog.fa.amb
for in1 in *clipped.1.fq.gz
do
	cd ../stacks_out/
	in2=${in2/.1./.2.}
	out=${in1/_R1_clipped.1.fq.gz/.sam}
	out2=${out/.sam/.sorted.bam}
	if [ ! -f ../alignments/$out2 ]
	then
		run_cmd.sh "bwa mem -t 3 catalog.fa $in1 $in2 > ../alignments/$out" ../alignments/$out
	fi
	cd ../alignments/
	run_cmd.sh "Convert_sam_to_sorted_bam.sh $out" $out2
done

cd ../alignments/
run_cmd.sh "samtools depth *.bam | egrep '^[0-9]+\s50' > coverage_repDepth.txt" coverage_repDepth.txt #this calculates coverage and pulls out the coverage for base 50 which should be representative of the contig
run_cmd.sh "bwa mem -t 3 $GENOME_LOCATION_GCF catalog.fasta > catalog_GCF.sam" catalog_GCF.sam
run_cmd.sh "awk '{if ( \$5 >= 50) { if ( \$2 == 0) { print \$1,\$2,\$3,\$4,\$5,\$6 } if (\$2==16){print \$1,\$2,\$3,\$4,\$5,\$6}}}' catalog_GCF.sam > catalog_GCF.partsam" catalog_GCF.partsam
run_cmd.sh "Annotate_AXY_Meriones_2.R" Autosomal.txt #now figure out the X and Y - especially for the tags in the vcf that have SNPs. 
run_cmd.sh "Get_Xlinked_tags.py -x Xlinked.txt -r ../stacks_out/rqtl_input_2.csvr -r2 ../stacks_out/rqtl_input_Var_2.csvr -o ../stacks_out/Xscaffs_for_rqtl.txt" ../stacks_out/Xscaffs_for_rqtl.txt
run_cmd.sh "recover_annotations.py -A Autosomal.txt -X Xlinked.txt -Y Ylinked.txt -U Unknown.txt -s catalog_GCF.sam -o Supplemental_File_4_Marker_annotation_and_scaffold.txt" Supplemental_File_4_Marker_annotation_and_scaffold.txt


#now will need to run: Meriones_Map_rqtl.R to make the map. final output file is Meriones_Map.csv
cd ../stacks_out/
if [ ! -f Meriones_Map.csv ]
then
	echo "Meriones_Map_rqtl.R needs to be run to make the Meriones_Map.csv. This takes a while and specific fine-tuning by hand and so is not hard-coded into this script. Run it now, create Meriones_Map.csv and then hit this script again."
	return  #this should stop the script - but I'm not 100% sure it will work (it didn't turn blue in bbedit), so I'll also throw in an exit. 
	exit
fi
#once Meriones_Map.csvr is made, then go ahead with this:


#need to re-make these with all the sites for hanging genes on the map
run_cmd.sh "make_rqtl_input_file.py -v ES_Fixed.recode.vcf -t \"_R1_clipped\" -p ../Phenotype_data.txt -o rqtl_input_all.csvr" rqtl_input_all.csvr # this takes only the first SNP per radtag to build the Map 
run_cmd.sh "rename_tags.py -s genotyped_radtags.sam -r rqtl_input_all.csvr > rqtl_input_2all.csvr" rqtl_input_2all.csvr #for hanging genes on the map later


	
#Then add on the het markers and the remaining hom markers:
run_cmd.sh "make_rqtl_input_file_for_Capture_hom.py -v ES_Fixed.recode.vcf -m Meriones_Map.csv -o hom_marker_dist_matrix.txt -c 20" hom_marker_dist_matrix.txt
run_cmd.sh "make_rqtl_input_file_for_Capture_het.py -v ES_Var.recode.vcf -m Meriones_Map.csv -o het_marker_dist_matrix.txt -c 20 -w PandF1.recode.vcf" het_marker_dist_matrix.txt
run_cmd.sh "evaluate_and_format_distance_matrix.R" evaluate_and_format_distance_matrix.R.finished
	

#Need to add het markers to the Map and re-annotate the RADtags with the proper genome (GENOME_LOCATION_GCF) - see line 52-56 above
#Grab het markers that are on the list called hetMarkers_placed.txt from csv: rqtl_input_Var_2.csvr and use the PandF1 dataset to modify the genotypes, then print them out and then add them to the Meriones Map 
#the add het markers script does actually add the hom markers too!
run_cmd.sh "cp Meriones_Map.csv Meriones_Map_AllMarkers.csv; add_het_markers_to_Meriones_Map.py >> Meriones_Map_AllMarkers.csv" Meriones_Map_AllMarkers.csv


#now run the RQTL on that file:
#this is really just to check that everything is in order - doesn't change much - map shrinks by about 35cM:
run_cmd.sh "Rscript ~/scripts/Meriones_Map_Het_markers_rqtl.R" Meriones_Map_AllMarkers_final.csv 


run_cmd.sh "get_details_of_mapping.py -g ${GENOME_LOCATION_GCF/.fna/.gff} -s ../alignments/catalog_GCF.sam -m Meriones_Map_AllMarkers_final.csv -o genes_on_map.txt 2>map_stats.txt" genes_on_map.txt
```
