## Supplemental_File_5_Codebase for "A high-density genetic map and molecular sex-typing assay for gerbils": recover_annotations.py

```
#!/usr/bin/env Python3


import sys
import re
import argparse
expl='''
recover_annotations.py
created by TDB 07-02-2019

Takes in the a list of markers on the X, Y, Auto, and Unknown, the alignments between the markers and the genome (GCF) and outputs some stats including a table of each marker, it's annotation, and it's scaffold, as well as a list of scaffolds that seems chimeric. 
'''
parser=argparse.ArgumentParser(description=expl)
parser.add_argument("--Autosomal", "-A", type=str, required=True, help="The Autosomal marker list.")
parser.add_argument("--Xlinked", "-X", type=str, required=True, help="The X-linked marker list.")
parser.add_argument("--Ylinked", "-Y", type=str, required=True, help="The Y-linked marker list.")
parser.add_argument("--Unknown", "-U", type=str, required=True, help="The unknown marker list.")
parser.add_argument("--sam", "-s", type=str, required=True, help="The .sam alignment of the catalog to the GCF genome with a sam header")
parser.add_argument("--map", "-m", type=str, required=True, help="The genetic map of the genome with all additional markers placed")
parser.add_argument("--out", "-o", type=str, required=True, help="Outfile name for the annotation list.")
args=parser.parse_args()


def parse_alignment(sam):
	print("parsing sam file:", sam, file=sys.stderr)
	with open (sam, "r") as SAM:
		for line in SAM:
			if not line.startswith("@"):
				bits=line.split("	")
				if int(bits[1]) != 2064 and int(bits[1]) != 2048: # ski markers that are multiply mapped
					if bits[0] not in the_dict:
						the_dict[bits[0]] = bits[2]
					
def parse_list(list, annotation):
	print("parsing list:", list, file=sys.stderr)
	with open(list, 'r') as LIST:
		for marker in LIST:
			marker = marker.strip("\n")
			if marker in map_dict:
				lg = map_dict[marker][0]
				cM = map_dict[marker][1]
			else:	
				lg = "NA"
				cM = "NA"
			print(marker, annotation, the_dict[marker], lg, cM, file=OUT, sep="	")

def parse_map(map):
	print("parsing map file:", map, file=sys.stderr)
	with open(map, "r") as MAP:
		for line in MAP:
			bits = line.split(",")
			if bits[0].startswith("NW"):
				rad = bits[0].split("_")[4]
			else:
				rad = bits[0].split("_")[0]
			lg = bits[1]
			cM = bits[2]
			if rad not in map_dict:
				map_dict[rad] = (lg, cM)
			else:
				if map_dict[rad][0] != lg:
					print("Error, rad", rad, "is on different LGs - probably has multiple SNPs. Needs to be removed from the map", file=sys.stderr)
	
the_dict = {}
map_dict = {}
with open(args.out, "w") as OUT:
	print("Marker", "Annotation", "Scaffold", "Linkage_Group", "centiMorgans", sep="	", file=OUT)
	parse_alignment(args.sam)
	parse_map(args.map)
	parse_list(args.Autosomal, "A")
	parse_list(args.Xlinked, "X")
	parse_list(args.Ylinked, "Y")
	parse_list(args.Unknown, "U")
```
