## Supplemental_File_5_Codebase for "A high-density genetic map and molecular sex-typing assay for gerbils": rename_tags.py

```
#!/usr/bin/env	Python3		
import sys
import argparse
import re
parser = argparse.ArgumentParser(description="written by TDB\n23-11-2017\nSpecifically for the Meriones_Map project.\nRenames the GBS tags so that they have the genomic scaffold attached to them.")                                              
parser.add_argument("-s", "--sam", type=str, required=True, help="A sam file of GBS tags alinged to the Meriones genome.", metavar="*.sam")
parser.add_argument("-r", "--rqtl", type=str, required=True, help="An rqtl file of GBS tags to rename.", metavar="*.sam")
parser.add_argument("--qual", '-q', type=int, required=False, default=50, help="minimum quality to assign a radtag to a scaf. Default: %(default)s")
parser.add_argument("--Out", "-o", type=str, required=False, help="Output file name. Omit for stdout")

args = parser.parse_args()
SAMFH = args.sam
rqtlFH = args.rqtl

if args.Out:
	OUT=open(args.Out, 'w')
else:
	OUT=sys.stdout	

pos_dict={}


def make_new_name(rad, snp):
	#print(rad, snp, pos_dict[rad])
	if rad not in pos_dict:
	#pos_dict[rad][0] =="*":
		new_name=rad+"_"+snp
	else:
		chr=pos_dict[rad][0]
		base=int(pos_dict[rad][1])+int(snp)
		rad_snp=rad+"_"+snp
		new_name=chr+"_"+str(base)+"_rad_"+rad_snp
		if chr not in scaff_placed:
			scaff_placed[chr]=0
		scaff_placed[chr]+=1		
	return(new_name)	


with open(SAMFH, 'r') as SAM:
	for line in SAM:
		if not line.startswith("@"):
			fields=line.split("	")
			rad=fields[0]
			chr=fields[2]
			start=fields[3]
			qual=int(fields[4])
			flag=int(fields[1])
			cigar=fields[5]
			matches=re.findall("(\d+M)", cigar)
			match_len=sum([int(i.strip("M")) for i in matches])
			#print(fields[0:4])
			#print(qual,flag,match_len, re.search("SA:Z:", line), file=sys.stderr)

			#if qual >= args.qual and (flag == 0 or flag == 16) and match_len > 200 and not re.search("SA:Z:", line):
				#print("\t\t",qual,flag,match_len, re.search("SA:Z:", line), file=sys.stderr)
			if not rad in pos_dict:
				pos_dict[rad]=(chr,start,qual)
	
	
				
	#for i in pos_dict:
	#	print(i, pos_dict[i])					
i=0	
scaff_placed={}			
with open(rqtlFH, 'r') as CSVR:
	for line in CSVR:
		i+=1
		if i>7:		 
			line=line.strip("\n")
			fields=line.split(",")
			
			if re.search("NW",line):
				cont_pos_rad_snp=fields[0]
				f=cont_pos_rad_snp.split("_")
				rad=f[3]
				snp=f[4]	
			elif re.search("NHTI", line):
				print("Error - you are using the wrong genome! The GCA one isn't annotated, use the GCF one!", file=sys.stderr)
				exit()
			else:
				rad_snp=fields[0]
				f=rad_snp.split("_")
				#print(f, file=sys.stderr)
				rad=f[0]
				snp=f[1]
			#if rad in pos_dict:
				#print(fields[0])
			new_name=make_new_name(rad, snp)
				#print(rad, pos_dict[rad], new_name)
			fields[0]=new_name
			print(",".join(fields), file=OUT)
		else:
			print(line, end="", file=OUT)		
			
print(len(scaff_placed),"unique scaffolds are associated with RADs with high confidence", file=sys.stderr)			
#for i in scaff_placed:
#	print(i, scaff_placed[i], )
```
