## Supplemental_File_5_Codebase for "A high-density genetic map and molecular sex-typing assay for gerbils": run_cmd.sh

```
#!/bin/bash
#this takes two inputs:
#1 the full command that you want to run in quote marks
#2 the name of the output file (not in quotes)
#this may mean that the output file is written twice; once in the command in quotes and once afterwards. 
cmd=$1
output=$2
echo "CMD: $cmd"
if [ ! -f $output ]
then
	echo "	evaluating"
	eval $cmd
fi
```
